# A simple genetic basis of adaptation to a novel thermal environment results in complex metabolic rewiring in Drosophila

**DOI:** 10.1101/174011

**Authors:** François Mallard, Viola Nolte, Ray Tobler, Martin Kapun, Christian Schlötterer

## Abstract

Population genetic theory predicts that rapid adaptation is largely driven by complex traits encoded by many loci of small effect. Because large effect loci are quickly fixed in natural populations, they should not contribute much to rapid adaptation. To investigate the genetic architecture of thermal adaptation - a highly complex trait - we performed experimental evolution on a natural *Drosophila simulans* population. Transcriptome and respiration measurements revealed extensive metabolic rewiring after only ∼60 generations in a hot environment. Analysis of genome-wide polymorphisms identified two interacting selection targets, *Sestrin* and *SNF4Aγ*, pointing to AMPK, a central metabolic switch, as a key factor for thermal adaptation. Our results demonstrate that large-effect loci segregating at intermediate allele frequencies can allow natural populations to rapidly respond to selection. Because *SNF4Aγ* also exhibits clinal variation in various *Drosophila* species, we suggest that this large effect polymorphism is maintained by temporal and spatial temperature variation in natural environments.

---

One of the major challenges in evolutionary genetics is to unravel the genetic architecture of phenotypic traits and how this affects the potential of natural populations to respond to selective forces. Theory predicts that evolution proceeds mainly through polygenic quantitative traits (Barton, Keightley 2002). Thus, adaptation is expected to involve rather subtle allele frequency changes at many small-effect loci (Le Corre, Kremer 2003; Pritchard, Pickrell, Coop 2010; Rockman 2012). Classic association studies as well recent whole genome association studies confirmed that variation at quantitative traits is due to a large number of loci and that large-effect loci are rare (Mackay, Stone, Ayroles 2009; Yang *et al.* 2010; Turner *et al.* 2011). Although, association studies are helpful to describe the genetic architecture of phenotypic traits, these genetic variants cannot be directly linked to adaptive responses (Franks, Hoffmann 2012). In contrast to these theoretical predictions, an increasing number of studies identified a small number of large effect loci which are driving rapid adaptation (reviewed in (Messer, Ellner, Hairston 2016)). Fluctuating selective pressure across time and space may contribute to the persistence of polymorphism at large-effect loci even over long evolutionary time scales (Savolainen, Lascoux, Merilä 2013; Lescak *et al.* 2015; Messer, Ellner, Hairston 2016). Importantly, these major effect loci typically encoded rather simple traits, such as melanism (Nachman, Hoekstra, D’Agostino 2003; vant Hof *et al.* 2016), insecticide resistance (Daborn *et al.* 2002) or lactose tolerance in humans (Bersaglieri *et al.* 2004). One noticeable exception is the evolution of song-less crickets, which occurred on two islands, but involved different major effect loci (Pascoal *et al.* 2014). It remains unclear to what extent rapid adaptation by large effect loci is an exception of simple traits, which are maintained in the population by fluctuating selection pressures

In the light of global warming, it is of key interest to understand how novel thermal environments can drive genetic adaptation and what is the nature of the associated phenotypic changes (Merilä, Hoffmann 2016; Scheffers *et al.* 2016). Temperature is a major abiotic factor known to affect a broad range of phenotypes and provides a good study system to investigate the genetic architecture of phenotypic evolution, in particular of quantitative traits. Most insight into thermal adaptation comes from contrasting natural populations that have evolved in different thermal habitats (Fabian *et al.* 2012; Bergland *et al.* 2014; Zhao *et al.* 2015; Machado *et al.* 2016; Sedghifar, Saelao, Begun 2016; Porcelli *et al.* 2016). Apart from complications intrinsic to natural populations, such as confounding signals of demography (Bergland *et al.* 2016) and the complexity of natural environments, the underlying evolutionary time scales are too long to be informative about rapid adaptation required to counter the current rate of climate change (Franks, Hoffmann 2012). The well-documented clinal variation and seasonal response of many genetic polymorphisms (Fabian *et al.* 2012; Bergland *et al.* 2014; Machado *et al.* 2016; Sedghifar, Saelao, Begun 2016) make *Drosophila* an excellent model system to study the impact of large-effect alleles segregating in natural populations on rapid adaptation to novel thermal environments. Shared clinal polymorphisms between two sister species, *D. melanogaster* and *D. simulans* suggest that genetic variants associated with thermal clines may have been segregating for a long evolutionary times.

Here we use experimental evolution in *D. simulans* to investigate the genetic architecture of phenotypic adaptation to a novel thermal environment. Transcriptomic data suggest that the evolved populations underwent a massive metabolic rewiring, which was confirmed by resting metabolism measurements. Whole genomic resequencing after ∼60 generations of evolution under our hot environment indicated that despite temperature adaptation being a complex trait, only a small number of selection targets were identified across 5 replicate populations. Two interacting loci associated with the AMPK, a key metabolic switch drive the phenotypic changes observed in our experiment. We show that these alleles are segregating at intermediate frequency in a European population and show a latitudinal cline in North American populations. These results suggest that experimental evolution identified variants, which play a key role in rapid spatial and temporal temperature adaptation in natural populations.

## Results

### Experimental evolution and phenotypic response

A natural *D. simulans* population from Póvoa de Varzim, Portugal was selected for ∼60 generations in a hot environment that fluctuated daily between 18 and 28°C (Figure 1). Five independently hot-evolved populations were compared to reconstituted ancestral populations and five populations that evolved in a cold environment fluctuating between 10 and 20°C. The cold-evolved control populations allowed us to rule out adaptation to culture conditions not specific to the hot environment (i.e. laboratory adaptation). To characterize the adaptive response of the evolved flies to high temperature, we assayed three phenotypes - fecundity, the whole transcriptome, and resting metabolism - in a common garden experiment at 23°C, the mean temperature of the experimental hot environment.

**Figure 1.**
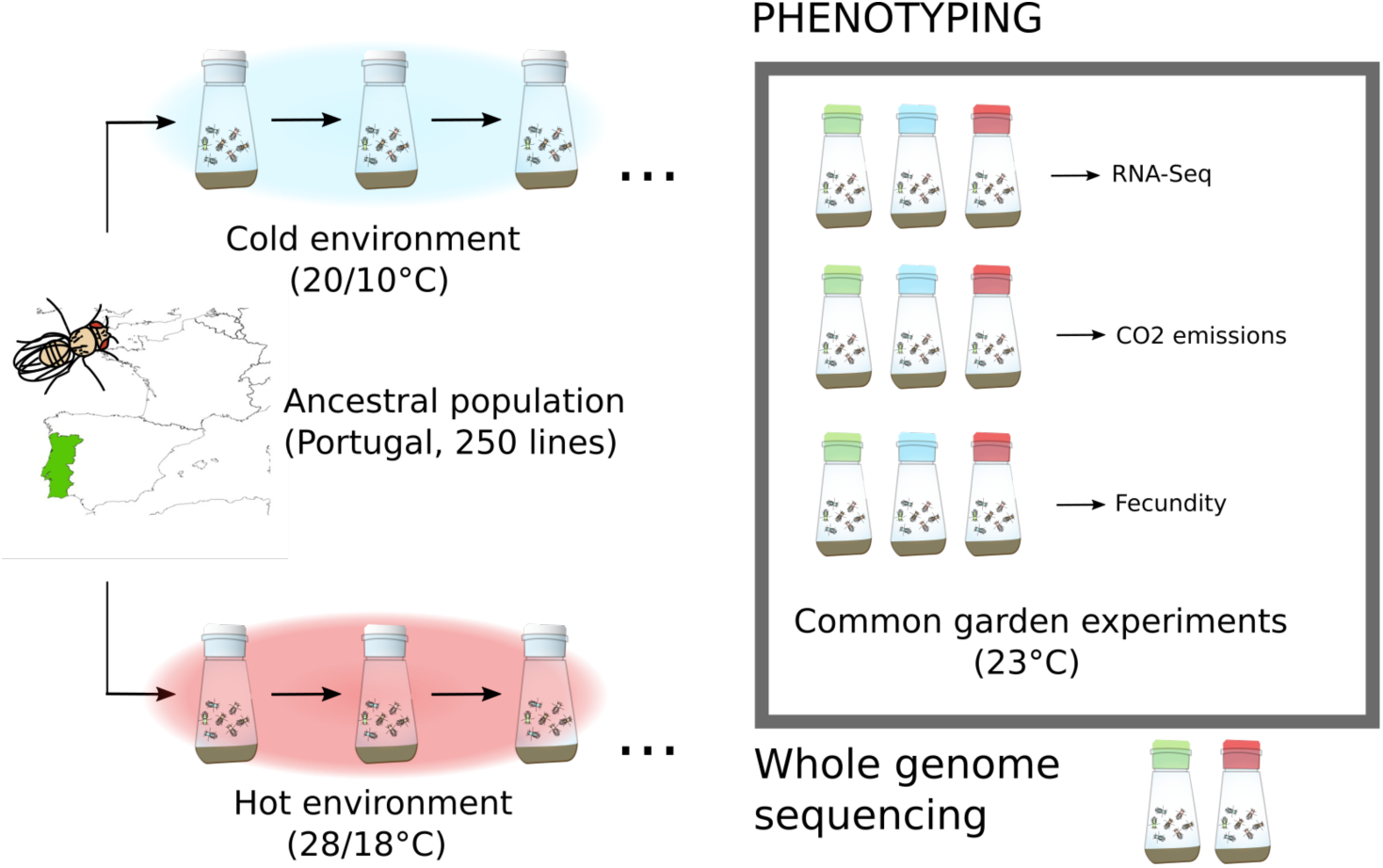
Experimental design. 250 isofemale lines from a natural *Drosophila simulans* population constituted the ancestral populations. The populations were kept at a constant populations size (1000 flies) with non-overlapping generations either in a hot environment (red) fluctuating between 28 and 18°C on a 12/12h cycle or a cold environment (blue), fluctuating between 20 and 10°C (five replicated population in each environment). We performed multiple phenotypic measurements on the evolved and reconstituted ancestral populations (green) in common garden experiments. We profiled the transcriptome of hot (F64) and cold (F39) evolved and reconstituted ancestral populations. Resting metabolism (i.e.: CO2 emission) and fecundity were measured later in the experiment (cold-evolved F74-F77; hot-evolved F127-F133). In addition we sequenced the entire genome of the ancestral and hot-evolved populations (F59). Figure supplement 1. Relative read coverage across genes used to control RNA-Seq libraries quality. Figure supplement 2. Mean normalized expression of chorion genes across RNA-Seq libraries used to investigate female contaminated samples.

Consistent with an adaptive response, the hot-evolved *D. simulans* populations were fitter than the ancestral population in the hot environment. In common garden experiments involving two different hot temperature regimes the hot evolved populations were more fecund (total number of eggs laid over successive 5 days) than the ancestral population (p=0.0006 and p<0.0001 at 23°C and 18/28°C cycling respectively, Figure 2 E, F). Similar to previous observations in *D. melanogaster* (Tobler, Hermisson, Schlötterer 2015), hot and cold evolved populations were only significantly different from each other at 23°C (p=0.0018, Figure 2 E), but not in the fluctuating temperature regime. These fitness differences suggest that the flies in our experiment adapted to a higher mean temperature as well as rapid temperature fluctuations (Tobler, Hermisson, Schlötterer 2015)

**Figure 2.**
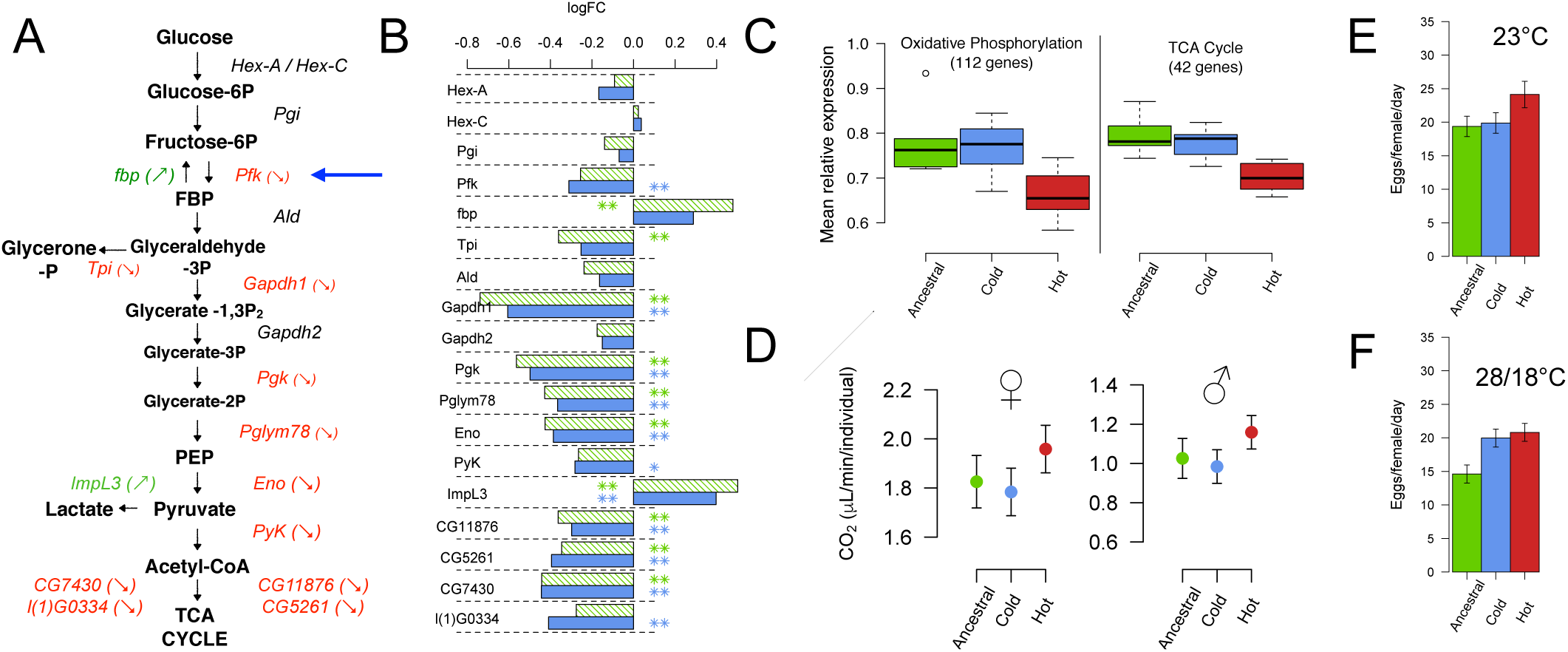
Phenotypic response of hot-evolved flies. (A) Evolution of gene expression in the glycolysis pathway. Enzymes significantly up (green) and down (red) regulated in the hot-evolved populations relative to the ancestral population are shown in color. (B) Gene expression changes (log_2_ fold change) in the hot-evolved populations relative to the ancestral (green) or cold-evolved (blue) populations. (** FDR<0.05, * FDR<0.1). (C) Hot-evolved populations (red) differ in gene expression from the ancestral (green) and cold-evolved (blue) populations for genes of the oxidative phosphorylation pathway and TCA cycle. (D) For both sexes, the resting metabolism of hot-evolved flies differs significantly from ancestral and cold-evolved populations. Both ancestral and cold-evolved populations are significantly different from the hot-evolved populations when considering both sexes in a single model (p=0.032 and p=0.003 respectively). Bars show mean ± 95% confidence intervals as estimated by our linear model (Material and Methods) (E, F) Higher fitness of hot-evolved flies: hot-evolved flies have a higher fecundity than the ancestral (p=0.0006) or cold-evolved populations (p=0.0018) at 23°C but differ only from the ancestral populations at 28/18°C (p<0.0001), which is consistent with previous results in *D. melanogaster*^13^ (see Supplementary Information for a detailed discussion). **Figure supplement 1.** Expression changes of fat metabolism and insulin signaling pathway genes **Figure supplement 2.** Additional resting metabolism measurements

We further characterized the molecular phenotype of the hot-evolved and control flies using RNA-Seq, collecting whole transcriptome data of young adult males (3-5 days old). Out of more than 9000 genes with reliable gene expression signals, 603 genes were differentially expressed (FDR<0.05) between hot-evolved and reconstituted ancestral populations. In contrast, the cold-evolved control populations were very similar to the ancestral ones, with only 41 genes differing significantly. Because 27 of these genes were also differentially expressed between ancestral and hot-evolved populations, we attributed them to non-temperature specific adaptation. The lack of statistically significant enrichment of functional categories among these 27 genes (Table S1) further substantiates that the majority of the transcriptional response observed in the hot-evolved flies is specific to adaptation to high temperatures.

Consistent with hot temperature adaptation affecting multiple genes, we identified a significant enrichment of several GO categories and KEGG pathways contrasting hot-evolved with ancestral or with cold-evolved populations. Genes down-regulated in the hot-evolved populations were enriched for more GO categories than up-regulated ones (Table S2). Up-regulated genes were mainly enriched for defense response (including Toll signaling pathway). Other categories overrepresented in up-regulated genes were triglyceride metabolism and cellular lipid metabolic processes, which include several genes involved in fatty acid synthesis or elongation (see Figure 2 – figure supplement 1). Down-regulated genes were enriched for a larger number of functions and pathways, most of which are related to metabolism: both TCA cycle and oxidative phosphorylation pathways were significantly down-regulated. Some key enzymes of glycolysis were also down-regulated (see Figure 2A,B) along with the sucrose metabolism and carbon metabolism pathways

### Resting metabolism measurements

With the transcriptomic data suggesting major regulatory changes affecting the metabolism, we reasoned that high-level metabolic phenotypes, such as respiration, should have changed as well. We quantified the resting metabolism by measuring CO_2_ emission overnight from the ancestral and both cold and hot evolved populations. After 127 generations in the hot and 74 generations in the cold environment we measured the CO_2_ emission of the evolved flies in parallel to a reconstituted ancestral population. In a GLM we identified the factors “population” (F_31,2_=5.7, P=0.008) and “sex” F_30,1_=13.7, P=0.0008) to have a significant effect on CO_2_ emission (see Fig 2E). “Body size” was not significant (F_30,1_=1, P=0.3), because the difference in weight between sexes was already explained by the factor “sex” and females produced significantly more CO_2_ than males.

We found similar results in a second series of measurements between the ancestral and hot evolved populations after 133 generations. Body weight was again non-significant (p=0.23), but the difference between hot evolved flies and the reconstituted ancestral population was only significant if body weight was included as a fixed effect in our model (see Figure 2 – figure supplement 2), suggesting that in addition to sex, other factors affected body size between populations. Since body size is an important factor influencing CO_2_ emission, we including body weight as a random effect in a mixed linear model and found a significant difference between the two populations (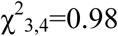, P=0.045). Because this value is close to the 5% threshold, we additionally tested for significance using bootstrapping by generating 10,000 random data sets having the same distribution as in our null model (only containing the random effect) and computed the test statistic for each of these data sets. The simulated test statistics are higher than the true one only 4.3% of the time, confirming the significant differences between the hot evolved and reconstituted ancestral populations.

### Genomic signature of adaptation

While the transcriptomic response is well suited to identify pathways that are altered in response to temperature adaptation, it is inadequate for pinpointing the causal mutation(s) driving these changes. To map the targets of selection, we performed Pool-Seq (Schlötterer *et al.* 2014) contrasts of the ancestral populations and hot-evolved flies at generation ∼60 to identify genomic regions harboring pronounced allele frequency changes across all five replicates. More than 2.7 million SNPs were tested for concordant allele frequency changes across replicates using the Cochran–Mantel–Haenszel (CMH) test. The Manhattan plot of the CMH –log10(p-values) showed a handful of pronounced peak structures (Figure 3A). Each peak comprises a set of linked SNPs that are highly differentiated between the hot-evolved and ancestral populations, a pattern indicating that the associated genomic regions likely carry selected variants. We estimated the effective populations size (*N*_e_) of 219 individuals (see Methods) in our populations. The 100 most significant SNPs mapped to 27 genes in *D. simulans* (false positive rate<0.04, Table S3). Two of these genes are involved in metabolism homeostasis and interact with each other: *Sestrin* and *SNF4Aγ* and are located on two distinct peaks on the 2R and 3R chromosome arms (red and blue arrows, Figure 3A).

**Figure 3.**
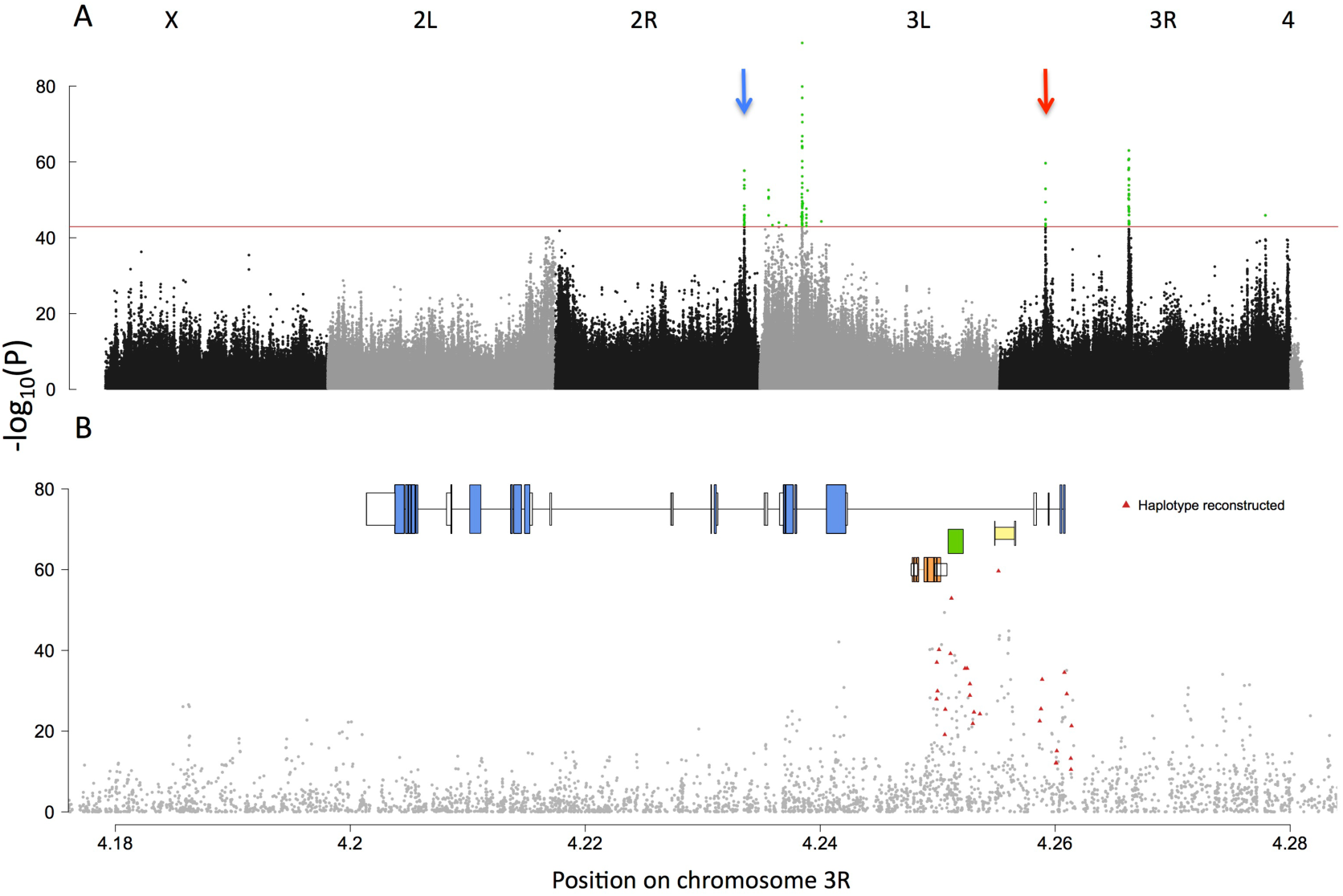
Genomic signature of adaptation to a hot laboratory environment. (A) Manhattan plot displaying the Cochran–Mantel–Haenszel p-values of 2,741,793 SNPs (Methods). The red line indicates the p-value cutoff for the most significant 100 SNPs (in green). The red and blue arrows indicate the *SNF4Aγ* and *Sestrin* peaks. (B) A close up of Manhattan plot around the *SNF4Aγ* region. On top of the Manhattan plot the gene structure of *SNF4Aγ* is shown together with three small genes (Snmp1, CG5810 and cDIP, see Table S3) located in one large intron of *SNF4Aγ*. Exons are indicated by colored boxes and introns by thin lines. White boxes indicate UTRs. Figure 3 – figure supplement 1. Allele frequency changes of the selected SNPs in the *SNF4Aγ* region. Figure 3 – figure supplement 2. Posterior distribution of the selection coefficient estimation in *SNF4Aγ* region. Figure 3 – figure supplement 3. A close up of the Manhattan plot around the *Sestrin* region. Figure 3 – figure supplement 4. Allele frequency changes of the selected SNPs in the *Sestrin* region. Figure 3 – figure supplement 5. Posterior distribution of the selection coefficient estimation in *Sestrin* region.

### Characterization of the *Sestrin* and *SNF4Aγ* loci

While the mapping precision of the genomic region containing *Sestrin* is not very high (see Supplementary Information), the selection signature for *SNF4Aγ* is narrow, encompassing less than 50 kb (Figure 3B). We further refined the selection signature by looking for correlated allele frequency trajectories across replicates (see Methods) and identified 27 SNPs that may reside on similar haplotypes in the ancestral population (Figure 3B). These 27 candidate SNPs start from a mean frequency of 44% in the ancestral population and rise as high as 96% (replicate 3) in ∼60 generations (mean increase 42%, Figure 3 – figure supplement 1). No selection signature was noticed in the *SNF4Aγ* or *Sestrin* regions in the cold-evolved control populations (data not shown). The posterior distribution of average selection coefficient for the 27 SNPs of interest in the *SNF4Aγ* locus results peaks around 0.06 (Figure 3 – figure supplement 2).

Around the *Sestrin* locus, we found a higher number of SNPs (96) distributed over a broader genomic region (∼60 kb) than for *SNF4Aγ* (See Figure 3 – figure supplement 3). The region that responded to selection contained multiple genes, *Sestrin* being at the left end of this region. Only the joint analysis of RNA-Seq and genomic data allowed the identification of the putative target of selection. The alleles of the selected haplotype start from a lower frequency than in the *SNF4Aγ* region (mean ∼20%, see Figure 3 – figure supplement 4) and increase by ∼40%. With the posterior distribution of the average selection coefficient peaking at 0.055 (Figure 3 – figure supplement 5), the selection strength of *Sestrin* was similar to *SNF4Aγ*.The broader genomic region around *Sestrin* can be explained by the lower starting frequency of the selected haplotype encompassing *Sestrin*.

### *SNF4Aγ* variants in the North American and Australian latitudinal clines

Because the SNPs, which responded to the new hot environment were present at intermediate frequency in the European population, we reasoned that they exhibit clinal variation in natural populations. We tested the 27 candidate SNPs in the *SNF4Aγ* region increasing in frequency in our experiment for clinal variation in natural populations from two different continents (Machado *et al.* 2016; Sedghifar, Saelao, Begun 2016). In the US populations described by Machado et al., we detected 20 out of the 27 candidate SNPs of the *SNF4Aγ* haplotype, which increased in frequency in the hot evolved populations. Eleven (55%) of these SNPs were at lower frequency in Maine than in Florida (see Figure 4 – figure supplement 1) and showed a clinal pattern among the Florida, Virginia and Maine populations (Figure 4), Samples from Pennsylvania, however, did not fit this clinal pattern and exhibited higher frequency than expected. From August to November, we observed a decrease in frequency of the hot alleles (i.e. those selected in the hot cages), an indication that our candidate SNPs could also be seasonal. This clinal frequency change is highly significant (p<0.008) based on Student’s t test for 10,000 random sets of SNPs generated by jackknifing 20 SNPs out of the 197 SNPs segregating in the region of interest (3R:4,249,000-4,261,000) (Figure 4 – figure supplement 2).

**Figure 4.**
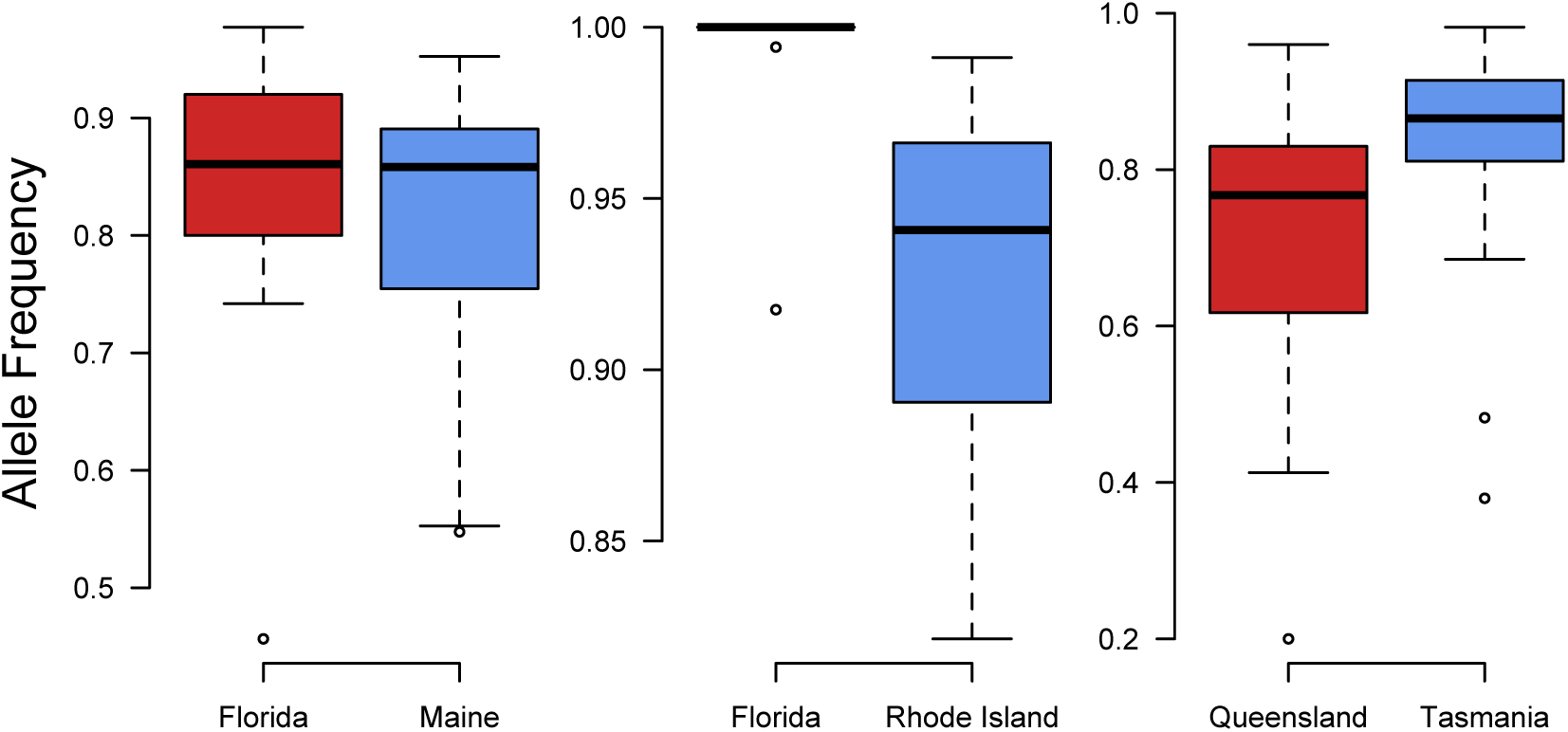
SNPs characterizing the selected haplotype at the *SNF4Aγ* locus in the experimental evolution study show clinal variation in natural populations. Allele frequencies of the alleles favored in the hot environment in: A. Machado et al. data set for Florida (red) and Maine (blue). 20 out of the 27 SNPs had sufficient coverage. B & C. Sedghifar et al. data set for B. North American cline (16 SNPs) and C. Australian cline (20 SNPs). Figure 4 – figure supplement 1. SNP based allele frequency differences along the North American (Machado et al. data set). Figure 4 – figure supplement 2. Distribution of t.statistics obtained from random sampling of SNPs in the region of interest for each of the clinal data sets.

In the Sedghifar et al. (Sedghifar, Saelao, Begun 2016) data set, we identified 16 and 25 SNPs from our list of 27 *SNF4Aγ* candidates in the North American and Australian populations, respectively. In the North American cline, we found a similar pattern as in the Machado et al. data set (Figure 4): in the Florida population, all the hot alleles were almost fixed (median allele frequency = 1) while they had lower frequencies in the Rhode Island population (median allele frequency = 0.94. This difference was significant based on the resampling test described above (Figure 4 – figure supplement 2). In the Australian cline, the pattern was inversed as the allele frequencies of our hot alleles were lower in the Queensland population (lower latitude, median = 0.77) than in the Tasmanian population (higher latitude, median = 0.87 (see Figure 4). Interestingly, this trend is consistent with the observation of Sedghifar et al. that clinal SNPs were preferentially going in the opposite direction.

## Discussion

### Novel thermal environment induces a rewiring of metabolic regulation

Temperature is a major factor modulating the expression of numerous genes in ectotherms and is particularly well studied in *Drosophila* (Zhao et al. 2015; Chen, Nolte, Schlötterer 2015; Porcelli et al. 2016). Our experimental populations, which evolved in a novel hot thermal environment displayed highly significant differences in gene expression involving many genes of well-defined pathways. Of particular interest were genes, which were down-regulated in the hot-evolved populations, because they suggest a global down-regulation of energy production in hot-evolved flies, affecting glycolysis, TCA cycle and oxidative phosphorylation pathways. Interestingly, a highly replicated study in *E. coli* found that RNA polymerase was the most frequently targeted gene across replicates, resulting in a lower rate of protein synthesis (Tenaillon *et al.* 2012), providing further evidence that an important evolutionary response to hot environments is to reduce the increase in energy production and protein synthesis, which is increased in hot environments and probably imposes a significant cost.

Consistent with modified metabolic rewiring of the hot-evolved populations, we found significant differences in CO_2_ production relative to the ancestral and cold-evolved control population (see Figure 2E & Figure 2 – figure supplement 2, see Supplementary Analysis). Contrary to naïve expectations, CO_2_ production was higher in the hot-evolved flies. Nevertheless, resting metabolism and gene expression are measured at two different moments of the daily cycle of the evolving populations, suggesting that the link between gene expression and energy production might not be straightforward. Additionally, higher CO_2_ production in hot-evolved flies is consistent with increasing O_2_ consumption associated with decreased AMPK activity (Johnson *et al.* 2010). Further insights into this counter-intuitive pattern of CO_2_ consumption comes from a metabolomic analysis of *D. melanogaster* under a wide range of developmental temperatures (Schou *et al.* 2017). At extreme temperatures the flies were depleted for sugars and energy metabolites (NAD+, NADP+ and AMP), which is attributed to their inability to maintain cellular homeostasis. If the hot conditions of our experiment have the same effect, flies not evolved to this environment may also be depleted for sugars and energy metabolites. In response, enzymes in the glycolysis, TCA cycle and oxidative phosphorylation pathways could be up-regulated. Hot evolved flies may have acquired the ability to maintain cellular homeostasis at high temperatures, allowing a higher resting metabolism without up-regulation of the metabolic pathway genes.

Our results contrast a recent study where CO_2_ production was conserved among *D. melanogaster* populations which evolved in different thermal environments (Alton *et al.* 2017). With several experimental details differing between the studies (isofemale lines vs. pools of outbred individuals, 20 minutes measurements during the day vs resting metabolism overnight) the interpretation of this apparent discrepancy is difficult. Nevertheless, it aligns well with the general controversy about the effect of temperature on the evolution of metabolism (Messamah *et al.* 2017). We conclude that the consistent differences in CO_2_ production between ancestral and evolved populations provide strong evidence of temperature specific evolution of metabolism regulation but also indicate that the underlying physiological changes are more complex.

## AMPK explains the phenotypic changes observed in hot evolved populations

Based on the genomic analyses alone, it is not possible to rule out other genes in the *Sestrin* peak as targets of selection, nor three other small genes that overlap with the selection signature of *SNF4Aγ* (Table S3). In combination with the expression data, however, the role of *SNF4Aγ* and *Sestrin* as the primary drivers of the metabolic rewiring becomes evident. *Sestrin* modulates the phosphorylation rate of the AMP-activated protein kinase (AMPK) (Ho *et al.* 2016)’ which is composed of *SNF4Aγ* and two other subunits. AMPK is a key player for energy homeostasis at Although the causal relationship is still unknown, we consider the robust differences in CO_2_ production to be a reliable readout of the metabolic rewiring in the hot-evolved populations. the cellular and the organismal levels, and both *SNF4Aγ* and *Sestrin* are directly linked to AMPK activity (Hardie 2007; Mihaylova, Shaw 2011; Ho *et al.* 2016). Low levels of ATP result in the activation of AMPK, which causes up-regulation of glycolysis and biogenesis of mitochondria (Reznick, Shulman 2006). Furthermore, energetically costly pathways, such as fatty acid production and gluconeogenesis are down-regulated by AMPK (Burkewitz, Zhang, Mair 2014). Inactivation of AMPK causes down-regulation of glycolysis and up-regulation of anabolic pathways such as fatty acid production, which were both seen in our data. Interestingly, *Pfk*, the target enzyme for AMPK in glycolysis, is the first down-regulated enzyme of the glycolysis pathway in our data set (Figure 2A, blue arrow). In *D melanogaster*, RNAi mediated down-regulation of *SNF4Aγ* increases glucose content of muscles and fat body (Ugrankar *et al.* 2015) and induces starvation behavior (Johnson *et al.* 2010). Some of the genes of the insulin receptor signaling pathway were also differentially expressed in the hot-evolved populations (Ilp6, InR, see Figure 2 – figure supplement 1). Moreover, some key enzymes involved in fatty acid production (ACCoAs, ACC and FASN2, Desat1, CG30008, CG33110, CG18609, see Figure 2 – figure supplement 1) show also some signal of up-regulation, consistent with the direct inhibition of ACC by AMPK (Hutber, Hardie, Winder 1997). Increased temperatures and heat stress deplete fat storage in *D. melanogast*er (Klepsatel *et al.* 2016) by invoking apoptosis in the fat body – a process dependent on *SNF4Aγ* (Lippai *et al.* 2008) that links the starvation-like expression pattern observed here to temperature adaptation. *Sestrin* is also connected with autophagy regulation in *Drosophila*, through its role in activating AMPK (Budanov, Karin 2008; Lee *et al.* 2010).

Thus, our results indicate that the activity of the key metabolic regulator AMPK is modulated through the differential regulation of the subunit *SNF4Aγ* and interacting gene *Sestrin* in hot-evolved populations. Given the central role of *SNF4Aγ* and *Sestrin* for temperature dependent metabolic rewiring, we reasoned that both genes should vary along temperature clines in natural populations. While we did not find evidence for clinality of *Sestrin*, the patterns for *SNF4Aγ* matched our expectations. A whole-genome polymorphism analysis identified *SNF4Aγ* as one of the top candidates in clinal North American *D. melanogaster* populations (Fabian *et al.* 2012). Clinal and seasonal variation of *SNF4Aγ* in *D. melanogaster* and *D. simulans* further implicate temperature as an adaptive driver (Bergland *et al.* 2014; Machado *et al.* 2016). Reanalyzing clinal population genetic data (Sedghifar, Saelao, Begun 2016), *SNF4Aγ* is among the 603 most differentiated genes shared by North American and Australian *D. simulans* populations. Gene expression of *SNF4Aγ* is clinal in European *D. subobscura* populations, with southern populations having lower expression levels (Porcelli *et al.* 2016), which parallels the response observed in our experimental evolution populations. Because the selected haplotype may be partially maintained in other populations, we tested the diagnostic SNPs for clinal variation. Remarkably, populations from the extreme ends of the North American cline exhibit a clinal signal for the diagnostic SNPs. Nevertheless, the signal was mixed for less extreme populations.

### Large-effect loci segregating at intermediate allele frequencies drive rapid evolution

The combined analysis of transcriptomic and whole genome resequencing data of a freshly collected *D. simulans* population evolving in a new thermal environment identified two genes, both connected to AMPK, a central metabolic switch. While many possibilities exist how metabolism could be regulated, the strong selection response in all replicates suggests that two major effect loci are driving the adaptive metabolic response in our populations. The observed selection signature clearly indicates that adaptation in our E&R study is dominated by a small number of loci with strong effect, providing another example for rapid adaptation driven by a few major effect loci (Daborn *et al.* 2002; Nachman, Hoekstra, D’Agostino 2003; Bersaglieri *et al.* 2004; Pascoal *et al.* 2014; vant Hof *et al.* 2016).

The two genes driving the metabolic switch in our experimental populations segregate at intermediate frequencies in the founder population and show clinal variation. Thus, it is highly plausible that these genes contribute to similar adaptive processes in natural populations, which probably occur over very short time scales. Because temperature varies seasonally, it is possible that spatial and temporal heterogeneity maintains the selected alleles at intermediate frequency in *D. simulans* (Savolainen, Lascoux, Merilä 2013; Bergland *et al.* 2014).

The fact that few large effect loci resulted in a clear selection signature in our experiment does, however, not preclude that several minor effect loci also influence the metabolic rewiring in hot environments. Previously, it had been shown that major effect alleles contributing to quantitative traits, show the fastest selection response, but with an increasing number of generations these loci are out-competed because small effect alleles gradually increase in frequency (Yeaman 2015). The reason for the loss of the large effect alleles is that it is easier to obtain genotypes close to the fitness optimum with small effect alleles, while large effect alleles could cause overshooting, resulting in more extreme phenotypes than favored by selection. Hence, the analysis of these experimental populations after a longer time-interval could be very informative to understand the dynamics of adaptive alleles in natural populations.

With the favored allele being fixed or close to fixation in southern populations in the US, it would be interesting to study the adaptive response in these populations. Because AMPK will probably not further contribute to adaptation, such an experiment could reveal other adaptive signals that were not detected in this study. Would such populations be segregating for other major alleles or would a polygenic response detected? We propose that E&R studies using different founder populations are a very powerful approach to answer questions about the genetic architecture of rapid adaptation, which are difficult to infer from natural populations.

### Materials and Methods

#### *Drosophila simulans* population sample

In summer 2008, we established 250 isofemale lines from a *Drosophila simulans* collection in Northern Portugal (Póvoa de Varzim). After about 10 generations in the laboratory, we generated 10 independent replicate ancestral. For each replicate we used five mated females from each of the 250 isofemale lines (1250 females in total). These females were distributed across five bottles containing 70 ml standard *Drosophila* medium.

#### Culture conditions during experimental evolution

The flies were propagated in two fluctuating temperature regimes: five replicates evolved in a hot treatment with 12 h at 18°C (dark) and 12 h at 28°C (light), the other five replicates in a cold treatment with 12 h at 10°C (dark) and 12 h at 20°C (light). The populations evolving in the two temperature treatments were processed in the same way, except that the time after which flies were transferred to a new bottle was adjusted to account for the slower development at lower temperatures. Approximately three days after eclosion of a new generation (four to five days in the cold treatment), all flies from the five bottles were combined. After careful mixing the adults within each replicate, five samples of 200 individuals each were transferred to fresh bottles. After 48h (72h) of egg-laying in the hot (cold) environment, adults were transferred again to fresh bottles for another 48h (72h), after which flies were either frozen or used for DNA extraction. To prevent selection for early fecundity, flies eclosing from the first transfer were only used if the second transfer did not yield enough flies to maintain a population size of 1000 individuals per replicate (5 bottles with 200 flies).

#### Common garden experiments

Prior to phenotypic assays and RNA-Seq, all five replicate populations of the hot and the cold evolved treatment as well as from a reconstituted ancestral population (Nouhaud *et al.* 2016) were maintained in a constant 23°C common garden environment for two generations to control for maternal effects.

At generation 64 in the hot and generation 39 in the cold environment the common garden experiments were set up from additional egg lays. These eggs were transferred to a constant temperature (23°C, 12:12h light/dark cycles). In parallel, an ancestral population was reconstituted from the isofemale lines as described in Tobler et al. 2015 (Tobler, Hermisson, Schlötterer 2015), a procedure, which faithfully mirrors the allele frequencies in the founder populations (Nouhaud *et al.* 2016). During the common garden experiment and phenotyping, this ancestral population was maintained in parallel to the evolved ones. After one generation of acclimatization, groups of 300 eggs were transferred to fresh bottles (i.e.: density control). Shortly after eclosion, adults were collected and sexed under CO2 anesthesia. Males were separated from females and recovered 24-36h from CO2 treatment before being frozen in liquid nitrogen at 2pm (approx. 6h after the light cycle started in the incubators).

#### Gene expression analysis

For all 15 populations (five replicates in the hot, cold and ancestral population) we generated two RNA-Seq libraries, each from different sets males. Total RNA was extracted from 25-30 males using the Qiagen RNeasy Universal Plus Mini protocol (Qiagen, Hilden, Germany) with DNase I treatment according to the manufacturer’s instructions. The RNA was quality-controlled on agarose gels and quantified using the Qubit RNA HS or BR Assay kit (Invitrogen, Carlsbad, CA). We generated strand-specific barcoded mRNA libraries using the NEBNext® Ultra Directional RNA Library Prep Kit for Illumina with a protocol modified to allow for a larger insert size than the default 200bp.

PolyA-mRNA was purified from 3μg total RNA and fragmented for 8 min. The 42°C incubation step in the first-strand synthesis and the 16°C step in the second-strand synthesis were extended to 30 and 90 min., respectively. Size selection for a target insert size of 330bp was performed using AMPure XP beads (Beckman Coulter, Carlsbad, CA). PCR enrichment was done using NEBNext Multiplex oligos following the recommended protocol with 12 PCR cycles and a 50 sec. extension step. The final libraries were bead-purified, quantified with the Qubit DNA HS Assay kit (Invitrogen, Carlsbad, CA) and pooled in equimolar amounts. Samples from ancestral, cold and hot evolved replicates were combined in the same pool, i.e. sequenced in the same lane, to reduce batch effects. Libraries were sequenced using a single-read 50bp protocol on a HiSeq2500. Two of these libraries had a too low coverage to be analyzed (Table S4) and we finally obtained data from 28 libraries (nine ancestral, nine cold and ten hot evolved).

#### RNA-Seq data processing and quality control

Raw reads were trimmed using PoPoolation (Kofler *et al.* 2011), quality threshold 20, minimum length 40) and aligned to the *Drosophila simulans* reference genome using GSNAP (Wu, Nacu 2010) using an hadoop cluster (Pandey, Schlötterer 2013). Throughout this study, we used the genome and annotation of Palmieri et al. 2015 (Palmieri *et al.* 2014) as default reference. All statistical analyses were performed using R (R Core Team 2016).

We performed several analyses to test the quality of each library. First we determined coverage heterogeneity, i.e.: a 3′ bias, using the geneBody_coverage tool implemented in the RSeQC package (Wang, Wang, Li 2012). Since the 3’ bias is most pronounced for long genes, we performed this analysis using the 20% longest genes of the *D. simulans* transcriptome (the bed file is available on demand). Strongly biased libraries were removed from all subsequent analyses (see Figure 1 – figure supplement 1 for an explanation of the cutoff used).

Based on 12 chorion and yolk protein genes we tested for female gene expression in male samples to identify female contamination due to sexing mistakes (Table S5). We excluded four outlier libraries that showed total log2 normalized expression of these genes higher than eight (see Figure 1 – figure supplement 2). After removing female contaminated libraries and libraries with 3′bias, a total of 20 libraries (five ancestral, seven cold and eight hot evolved libraries) remained for analysis. Spearman correlation coefficients between all libraries showed a correlation of at least 0.948 between libraries. Despite some heterogeneity among libraries, a multi-dimensional scaling plot did not show outliers. Therefore, we retained all 20 remaining samples for differential gene expression analysis.

Read counts were determined with Rsubread (Liao, Smyth, Shi 2013) and differentially expressed genes were identified with the EdgeR (Robinson, McCarthy, Smyth 2010; Schurch *et al.* 2016). We normalized gene expression levels with the TMM method, restricting our analysis to the 70% most highly expressed genes (minimum count per million (CPM): 6.23, 9238 genes). We used negative binomial GLMs to estimate the effect of selection regime on gene expression. We then computed ad hoc contrasts to find differentially expressed genes between groups of interest. The Benjamini-Hochberg procedure was applied to control for false discovery rate (FDR < 0.05).

Using these differentially expressed genes, we performed KEGG pathway enrichments with the R package “gage” (Luo *et al.* 2009) using the logFC computed by EdgeR. GO enrichment analyses were performed with Gorilla (Eden *et al.* 2009), where all genes retained after filtering were used as the background data set. We only considered genes differentially expressed in the comparison of the hot samples against the ancestral as well as against the cold populations. Furthermore, in the comparison ancestral against the cold population the |logFC| was smaller than 0.2. (reported in Table S2)

#### Pool-Seq analysis

Genomic DNA for pooled sequencing of the ancestral and the hot evolved flies from generation F59 was extracted from females only. For each evolved population DNA was extracted from about 500 females. Since only two replicates of the founder females were frozen (1250 females each), we added one replicate from generation F2 as a substitute. Genomic DNA was extracted using either the DNeasy Blood and Tissue Kit (Qiagen, Hilden, Germany) (for the two ancestral replicates) or a high salt extraction protocol (Miller, Dykes, Polesky 1988) including RNase A treatment for the evolved populations. Paired-end libraries were prepared with different protocols and sequenced on different Illumina platforms (see Table S6 for details). When the coverage of the first runs was not sufficient, we re-sequenced the populations at a later stage with a HiSeq2500 in 2x120bp runs which required a modified adapter configuration (see Table S6). The reads of the ancestral and the generation F2 replicates were combined and then randomly split into five artificial data sets to serve as replicated ancestral populations that match the number of replicates in the hot evolved populations. SNPs were called with PoPoolation2 keeping only sites with at least one read with the minor allele per sample and coverage between 5 and 500. We masked sites flanking indels (± 5bp) and repeats using PoPoolation2 (Kofler, Pandey, Schlötterer 2011) and RepeatMasker (www.repeatmasker.org, file available on demand). While library preparation protocols do not affect allele frequency estimates based on Pool-Seq (Kofler, Nolte, Schlötterer 2016), insert size does (Kofler *et al.* 2016). Therefore, we mapped the trimmed reads (quality ≥ 20 and length ≥ 50 bp) to the reference genome (Palmieri *et al.* 2014) using three different mappers (bowtie2 (Langmead, Salzberg 2012), bwa-mem (Li, Durbin 2009) and NOVOALIGN (Novocraft 2010). Using only the SNPs called with all three mappers and restricting the analysis of each SNP to the mapper resulting in the least significant comparison (CMH test), prevents false positives (Kofler *et al.* 2016). We filtered for proper pairs and mapping quality ≥ 20 and finally retained 2,741,793 SNPs. We used Wright-Fischer simulations to estimate allele frequency changes expected in the absence of selection. Five independent simulation runs were performed, matching the initial frequency distribution of our ancestral populations and the number of SNPs tested in the original data set. We then added sampling noise to mimic the Pool-seq process (binomial sampling) and conducted CMH tests on these simulated data, similarly as for the empirical dataset. This way, we obtained a null distribution of CMH-based p-values under a null hypothesis. Given a certain p-value threshold, the false positive rate was computed as the fraction of simulated (neutral) and empirical loci. All simulations, adding sampling noise and performing CMH-tests were done with the poolSeq R-package (Taus et al. *in prep*).

27 genes contained at least one candidate SNP (Table S3). Since *SNF4Aγ* was a good candidate to explain the observed phenotypic changes, we focused the subsequent analysis on a 200kb region on chromosome 3R (4,150,000:4,350,000, see Figure 2B). Reasoning that candidate SNPs may be located on one or a few haplotypes only, we used a haplotype reconstruction method that relies on the identification of SNP markers showing a correlated response across replicates and time points (Franssen, Barton, Schlötterer 2016). Allele frequencies for this analysis were based on NOVOALIGN. Using the software package haploReconstruct (Franssen, Barton, Schlötterer 2016) we identified SNPs with a correlation of least 0.95. We found 27 diagnostic SNPs (minimum coverage 20, minimum allele frequency change 0.2 in all 5 replicates).

We estimated the effective population size (Ne) using the method of Jonas et al. (Jónás *et al.* 2016), which accounts for the sampling procedure of Pool-Seq (plan I) based on all polymorphic sites of the chromosomes 2 and 3. We used WFABC (Foll, Shim, Jensen 2015) to infer the selection coefficient (s) based on the N_e_ estimate of 219 and mean allele frequency of the 27 SNPs across all three replicates at the start of the experiment and at generation 59. Since WFABC cannot deal with replicates, we calculated a consensus allele frequency trajectory for each of our 27 SNPs across all 5 replicates. We averaged the posterior distribution of each SNP to obtain a consensus distribution (Figure 3 – figure supplement 2). The same procedure was then repeated for the *Sestrin* locus (2R-17520000:17600000, see Figure 3 – figure supplement 3). Using the haplotype reconstruction method, we found 96 diagnostic SNPs (minimum coverage 20, minimum allele frequency change 0.2 in all 5 replicates) showing correlated allele frequency changes (Figure 3 – figure supplement 4). We estimated the selection coefficient (s) based on these 96 SNPs (Figure 3 – figure supplement 5) using the same Ne estimate (219).

#### Resting metabolism

Resting metabolism was determined by repeatedly measuring overnight CO_2_ emission using a stop-flow respirometry system (Sable Systems International). All replicates of evolved and ancestral populations were reared for two generations at 23°C, controlling egg density (400 eggs per bottle). Flies were collected shortly after eclosion and after 24h males and females were separated and placed at low density in vials (25 flies/cm3) under CO2 anesthesia. After 48h recovery, the CO2 emission of 3-5 days old males and females was measured. Each assay was conducted at 23°C in the dark, overnight (at least 12h). The flies from different replicates were randomly assigned to one out of 8 chambers together with a small piece of fly food (2 cm3) to avoid starvation response and desiccation. During the assays, a multiplexer (RM8 Intelligent Multiplexer) is sequentially flushing the metabolic chambers. Each flushing cycle lasts 5 minutes at a constant flow rate of 50μl/min. We obtained repeated measurements in 40 minutes intervals for each of the eight channels. After removal of water by passing through a Magnesium perchlorate column, we measured CO_2_ with CA-10A Carbon dioxide Analyzer. For each flushing cycle we determined the total CO_2_ emission using the ExpeData software (Sable Systems International) with an in house script (available on demand). At the end of each assay the flies were dried and weighted.

We conducted the assay after generation 127 contrasting all three populations (2 ancestral populations, 4 cold and 4 hot evolved populations, 5 successive runs), and a second time after 133 generations (5 hot replicates and 4 ancestral replicates, 4 runs). We estimated the resting metabolism for each chamber as the average of the three lowest observations overnight (Jensen *et al.* 2014). During the first set of measurements (generation 127), one chamber was left empty as a negative control. Because the CO_2_ levels in the empty chamber were always very low compared to the CO_2_ levels of chambers containing flies, we used all 8 chambers with flies for the second set of measurements (generation 133). We analyzed the data with linear models including mean dried weight and population identity (ancestral, cold or hot evolved) as fixed effects. Significance of the differences between populations was tested using ANOVA F-tests. Assumptions of the models (normality of the residuals and homogeneity of the variance) were validated by visual inspection of the residuals.

#### Fecundity assays

We conducted fecundity assays of the cold (F78) and hot (F133) evolved populations in parallel with a reconstituted ancestral population using the same common garden design as described above. The only modifications were that we performed two generations of density control rather than a single one and that the ancestral population was reconstituted from ∼90-100 lines only. In parallel, we set up another common garden experiment that differed only in the temperature regime, which was not constant but cycled between 28/18°C light/dark conditions.

Flies were collected shortly after eclosion and allowed to mate within a 24h period. Approximately 60 flies were placed under CO_2_ anesthesia in separate bottles to estimate egg laying rates. We created two bottles for each replicate (30 bottles in total). Every day, all flies were transferred without CO_2_ anesthesia to fresh food and the eggs laid were counted. We did not count the eggs laid during the first 24 hours after anesthesia and recorded the next 5 consecutive days. After six days the flies were then sexed and counted. We determined the mean number of eggs per female in a 24h interval for each replicated population and tested for differences between populations using linear models in R. Significance of the fixed effects was tested using ANOVA F-tests and differences between populations using Tukey tests (using the multcomp library and appropriate contrasts). At 23°C, we excluded the results of two hot evolved replicates (the 3rd and 4th) due to problems with the density control and the data from only three replicates were used.

##### Reanalysis of latitudinal North American and Australian *D. simulans* populations

We analyzed two published data sets of *D. simulans* populations along a cline (Machado *et al.* 2016; Sedghifar, Saelao, Begun 2016). First, we used *F*_ST_ to determine whether the *SNF4Aγ* and *Sestrin* regions were differentiated along the clines. Since *F*_ST_ values do not indicate the direction of allele frequency differences, we performed a second analysis testing specifically whether the difference in allele frequencies between Northern and Southern populations along a cline differs in the direction expected based on our experiment. Below, we describe the analysis for each of the two data sets separately.

We downloaded, trimmed and mapped the raw data from Machado et al. (Machado *et al.* 2016) using the same pipeline as described above but using only a single mapper (NOVOALIGN). All alignment files were converted into an mpileup file using samtools. We restricted our SNP based analysis to the individual sequences that were collected from four populations from Florida, Virginia, Pennsylvania and Maine (thus excluding the pools). The Pennsylvania population was sampled three times the same year in August, September and November 2011. We called the 27 SNPs of *SNF4Aγ* occurring on a haplotype block in the Portugal population in each population. At all 21 positions, we classified each sample in four categories: homozygous for either allele, heterozygous or unknown. As the coverage for each sample was relatively low (1.8-fold), all reads were used to determine the genotype of each individual. Based on this classification, we computed the frequency of all the rising *SNF4Aγ* variants along the cline for each position. Only SNPs were used for which we genotyped in at least half of the samples in each population, which reduced the number of SNPs to 20.

We downloaded, trimmed and mapped the raw data from Sedghifar et al. (Sedghifar, Saelao, Begun 2016) using the same pipeline as described above but using only a single mapper (NOVOALIGN). All alignment files were converted into an mpileup file using samtools and we computed *F*_ST_ between populations at each polymorphic position using PoPoolation2 (Kofler, Pandey, Schlötterer 2011). We filtered for TE insertions, sites flanking indels (± 5bp) and only retained SNPs with a minimum coverage 10. The *F*_ST_ values were computed for all positions individually while accounting for different numbers of chromosomes in each pool (6,795,806 and 7,990,580 SNPs in the US and Australian clines respectively). Focusing on the SNPs with the strongest differentiation, we retained only the top 0.1% SNPs with the highest *F*_ST_ values, resulting in *F*_ST_ thresholds of 0.53 and 0.52 in the Australian and American populations, respectively. These SNPs mapped to 2053 and 1506 genes, out of which only 603 genes were shared. Additionally, we compared the allele frequencies of our SNPs of interest in the *SNF4Aγ* region across the two clines (*Sestrin* was not among the 603 shared genes). We called the SNPs on each cline separately and only retained SNPs with a minimum coverage of 20 in each of the sampled sites.

## ACKNOWLEDGMENTS

This work was supported by a Marie Skłodowska Curie Individual Fellowship (H2020-MSCA-IF-661149) to F.M. and the European Research Council (ERC) grant “ArchAdapt” awarded to C.S and the Austrian Science Funds (FWF, W1225). We are thankful to Pablo Orozco-terWengel for fly collection and to the members of the Institut für Populationsgenetik for feedback and discussion. Some Illumina sequencing was performed at the VBCF NGS Unit (<u>www.vbcf.ac.at</u>). The authors declare that they have no competing interests.

---

**Table S1.** List of genes consistently down- or up-regulated in the contrast between the ancestral and the two evolved populations (laboratory adaptation). The table presents the log2 fold changes and p-values for each comparison ancestral-cold evolved and ancestral-hot evolved. Negative log2 fold changes indicate that the expression is decreased in the evolved populations (and reciprocally).

| FBgn | Symbol | log2FC (Anc/Hot) | P (Anc/Hot) | log2FC (Anc/Cold) |
| --- | --- | --- | --- | --- |
| FBgn0034612 | CG10505 | -1.904062224 | 2.29645E-16 | -1.015207739 |
| FBgn0030332 | CG9360 | -1.490895749 | 3.34E-15 | -1.200243418 |
| FBgn0032381 | Mal-B1 | -1.040488713 | 2.90E-06 | -1.322138651 |
| FBgn0000473 | Cyp6a2 | -0.993236165 | 7.07E-06 | -1.244721364 |
| FBgn0033204 | CG2065 | -0.926040147 | 7.97E-10 | -0.636353128 |
| FBgn0033065 | Cyp6w1 | -0.881128669 | 2.83E-04 | -1.314932308 |
| FBgn0033981 | Cyp6a21 | -0.774379264 | 2.16E-10 | -0.510986228 |
| dsim_PG00121 | - | -0.754138774 | 8.02E-08 | -0.876602722 |
| FBgn0013773 | Cyp6a22 | -0.717022506 | 2.12E-05 | -0.698016808 |
| FBgn0027600 | obst-B | -0.701498698 | 1.43E-07 | -0.523207216 |
| FBgn0000008 | a | -0.696467292 | 1.29E-07 | -0.541119809 |
| FBgn0038734 | CG11453 | -0.600415372 | 4.23E-06 | -0.554513603 |
| FBgn0038516 | P5cr-2 | -0.520118726 | 1.88E-05 | -0.501495527 |
| FBgn0030615 | Cyp4s3 | -0.417536345 | 3.77E-05 | -0.456748501 |
| FBgn0033888 | CG18568 | -0.286296192 | 1.33E-03 | -0.380724409 |
| FBgn0261800 | LanB1 | 0.305136437 | 0.002419601 | 0.40196194 |
| FBgn0037000 | ZnT77C | 0.383888163 | 5.47E-05 | 0.397886895 |
| FBgn0259140 | CG42255 | 0.457389454 | 4.41E-04 | 0.547324689 |
| FBgn0265267 | CG18258 | 0.701005005 | 1.57319E-10 | 0.430466065 |
| FBgn0032136 | Apoltp | 0.765447259 | 2.68E-14 | 0.449187463 |
| dsim_PG00244 | - | 1.004387037 | 3.13E-04 | 1.540294833 |
| FBgn0032322 | CG16743 | 1.054180333 | 7.73E-12 | 0.596855797 |
| FBgn0035619 | CG10592 | 1.284373376 | 1.61E-09 | 0.803228768 |
| FBgn0051324 | CG31324 | 1.413172663 | 2.41E-20 | 0.653213722 |
| FBgn0033729 | Cpr49Af | 1.591154299 | 1.52E-03 | 1.938187162 |
| FBgn0035620 | CG5150 | 1.610327819 | 2.20E-14 | 0.83708478 |
| FBgn0039471 | CG6295 | 1.741706632 | 3.75995E-09 | 1.272948225 |

**Table S2**

Results of the gene ontology enrichment and Kegg-Pathway analysis. Sheet 1: final list of genes used for the analysis, filtered as described in the Materials and Methods section. Sheet 2&3: up and down-regulated GO categories classified by process, component and function. Sheet 4: Kegg Pathways differentially expressed between ancestral or cold and hot evolved populations.

**Table S3**

List of the 27 genes containing at least one SNP among the 100 SNPs with the most extreme p-values from our CMH test.

**Table S4**

Number of mapped reads for the sequenced data. Genomic data: total number of mapped reads after all filtering steps before any analysis for each mapper separately. RNA-Seq: total number of mapped reads and counted reads after all filtering steps.

**Table S5**

Count per million (after TMM normalization) of each of the 12 female biased genes used to isolate female contaminated RNA-Seq libraries.

**Table S6.**
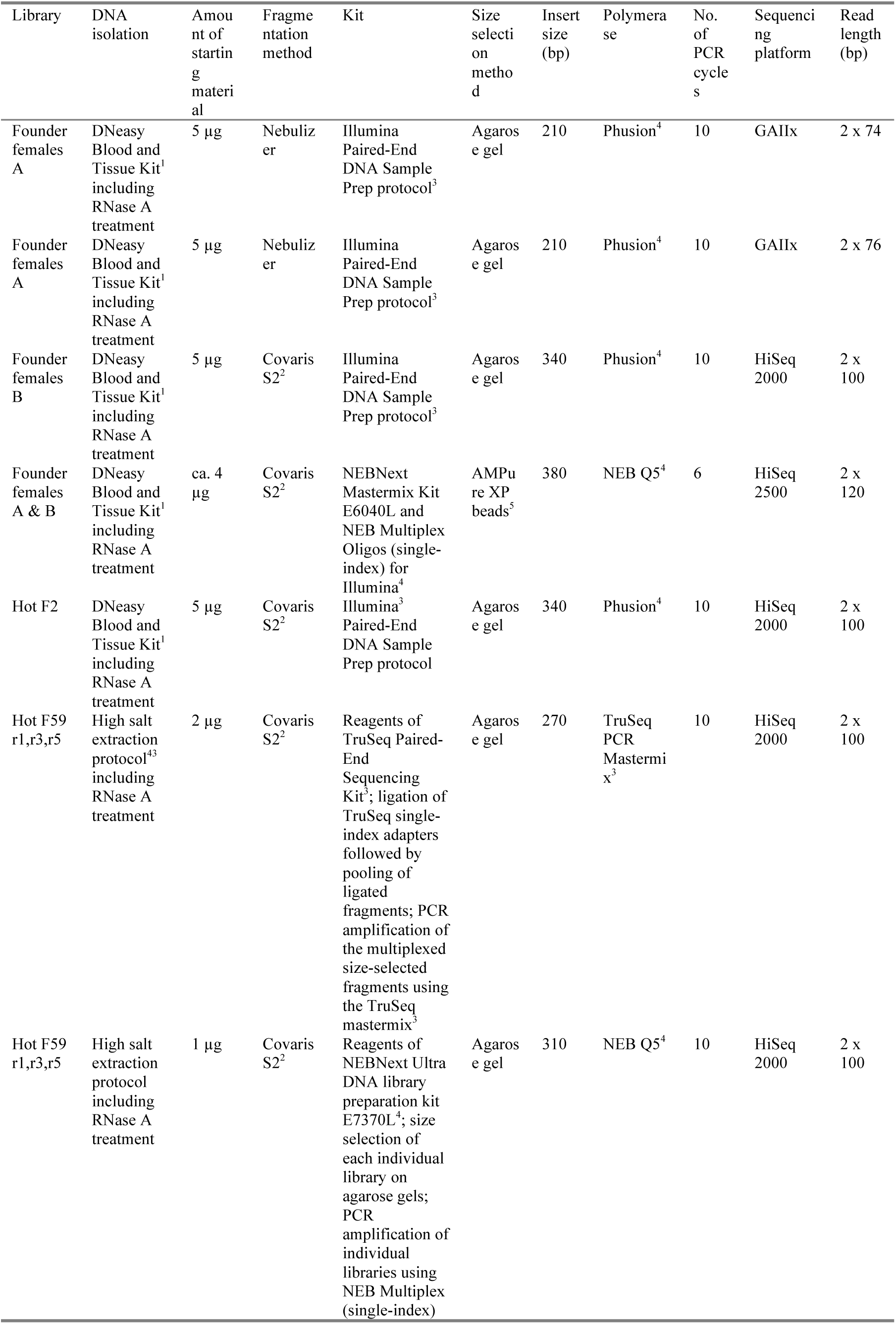
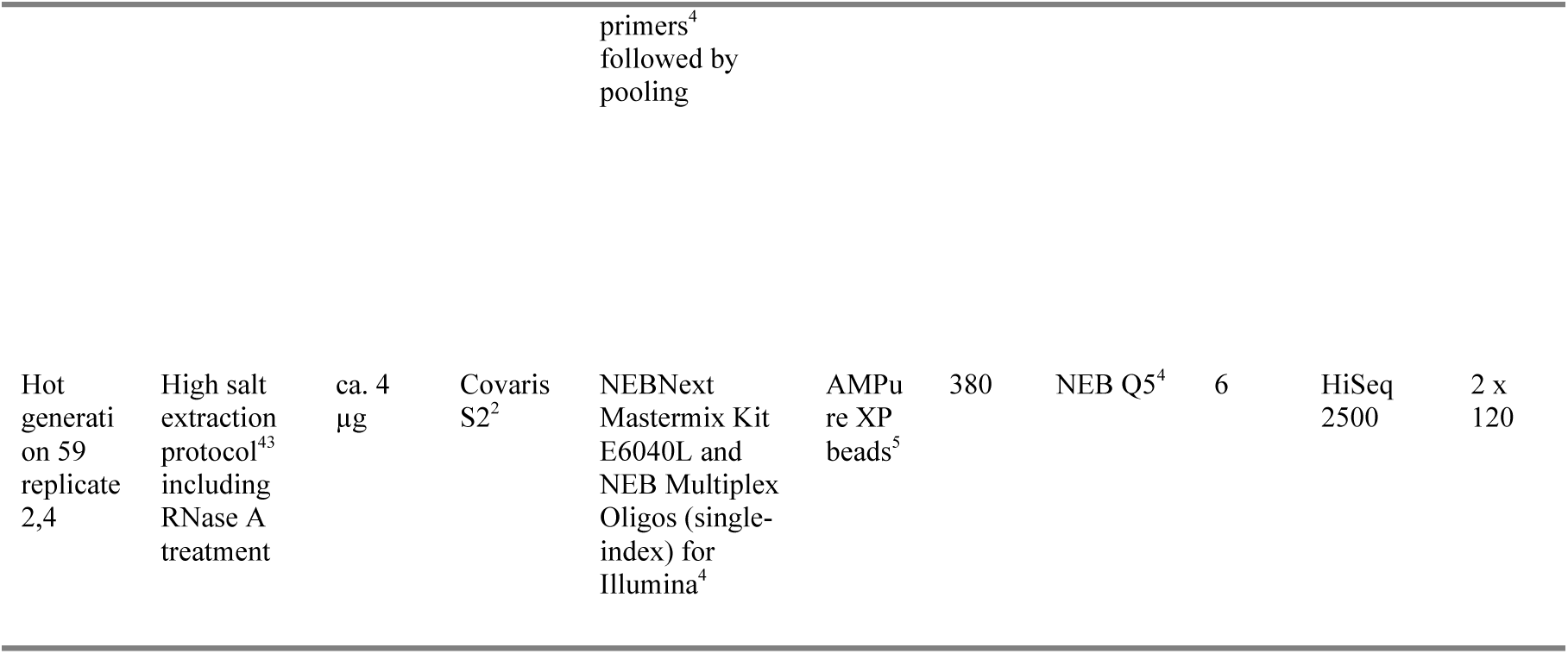
Description of the extraction and library preparation methods used for each sample (genomic data).

| Library | DNA isolation | Amount of starting material | Fragmentation method | Kit | Size selection method | Insert size (bp) | Polymerase | No. of PCR cycles | Sequencing platform | Read length (bp) |
| --- | --- | --- | --- | --- | --- | --- | --- | --- | --- | --- |
| Founder females A | DNeasy Blood and Tissue Kit<sup>1</sup> including RNase A treatment | 5 µg | Nebulizer | Illumina Paired-End DNA Sample Prep protocol<sup>3</sup> | Agarose gel | 210 | Phusion<sup>4</sup> | 10 | GAIIx | 2 x 74 |
| Founder females A | DNeasy Blood and Tissue Kit<sup>1</sup> including RNase A treatment | 5 µg | Nebulizer | Illumina Paired-End DNA Sample Prep protocol<sup>3</sup> | Agarose gel | 210 | Phusion<sup>4</sup> | 10 | GAIIx | 2 x 76 |
| Founder females B | DNeasy Blood and Tissue Kit<sup>1</sup> including RNase A treatment | 5 µg | Covaris S2<sup>2</sup> | Illumina Paired-End DNA Sample Prep protocol<sup>3</sup> | Agarose gel | 340 | Phusion<sup>4</sup> | 10 | HiSeq 2000 | 2 x 100 |
| Founder females A & B | DNeasy Blood and Tissue Kit<sup>1</sup> including RNase A treatment | ca. 4 µg | Covaris S2<sup>2</sup> | NEBNext Mastermix Kit E6040L and NEB Multiplex Oligos (single-index) for Illumina<sup>4</sup> | AMPure XP beads<sup>5</sup> | 380 | NEB Q5<sup>4</sup> | 6 | HiSeq 2500 | 2 x 120 |
| Hot F2 | DNeasy Blood and Tissue Kit<sup>1</sup> including RNase A treatment | 5 µg | Covaris S2<sup>2</sup> | Illumina<sup>3</sup> Paired-End DNA Sample Prep protocol | Agarose gel | 340 | Phusion<sup>4</sup> | 10 | HiSeq 2000 | 2 x 100 |
| Hot F59 r1,r3,r5 | High salt extraction protocol<sup>43</sup> including RNase A treatment | 2 µg | Covaris S2<sup>2</sup> | Reagents of TruSeq Paired-End Sequencing Kit<sup>3</sup>; ligation of TruSeq single-index adapters followed by pooling of ligated fragments; PCR amplification of the multiplexed size-selected fragments using the TruSeq mastermix<sup>3</sup> | Agarose gel | 270 | TruSeq PCR Mastermix<sup>3</sup> | 10 | HiSeq 2000 | 2 x 100 |
| Hot F59 r1,r3,r5 | High salt extraction protocol including RNase A treatment | 1 µg | Covaris S2<sup>2</sup> | Reagents of NEBNext Ultra DNA library preparation kit E7370L<sup>4</sup>; size selection of each individual library on agarose gels; PCR amplification of individual libraries using NEB Multiplex (single-index) | Agarose gel | 310 | NEB Q5<sup>4</sup> | 10 | HiSeq 2000 | 2 x 100 |
|  |  |  |  | primers<sup>4</sup><br>followed by<br>pooling |  |  |  |  |  |  |
| Hot<br>generati<br>on 59<br>replicate<br>2,4 | High salt<br>extraction<br>protocol<sup>43</sup><br>including<br>RNase A<br>treatment | ca. 4<br>µg | Covaris<br>S2<sup>2</sup> | NEBNext<br>Mastermix Kit<br>E6040L and<br>NEB Multiplex<br>Oligos (single-index) for<br>Illumina<sup>4</sup> | AMPu<br>re XP<br>beads<sup>5</sup> | 380 | NEB Q5<sup>4</sup> | 6 | HiSeq<br>2500 | 2 x<br>120 |

**Figure 1 – figure supplement 1.**
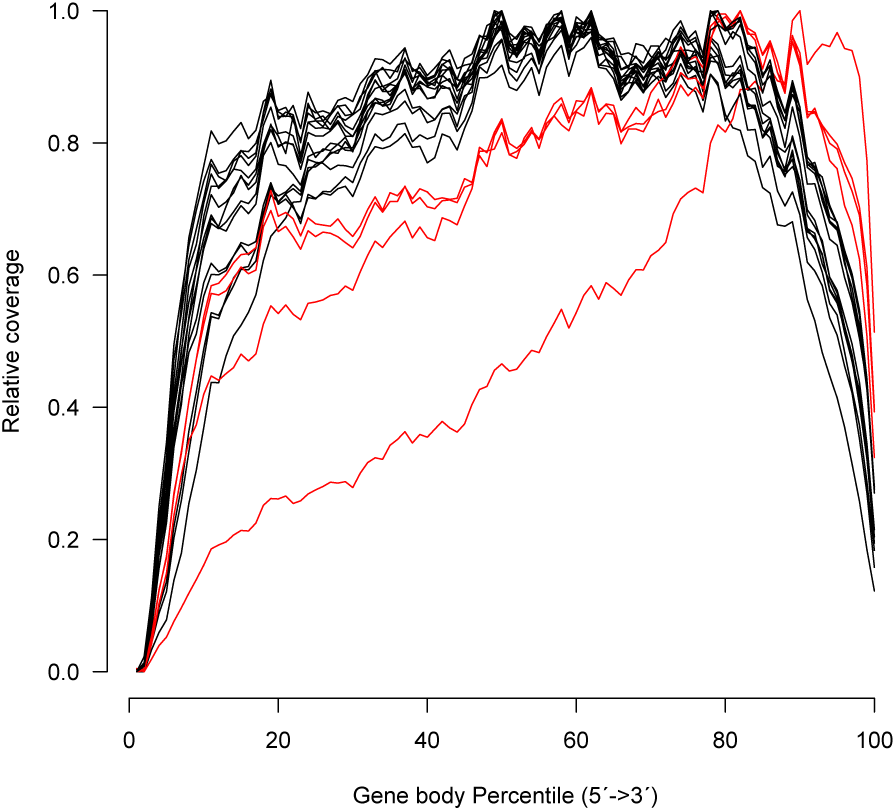
Relative coverage across genes for 28 RNA-Seq libraries. We removed three libraries (in red) from our analysis, because they exhibited a very pronounced 3’ bias: at 50% gene body length the relative coverage was below 90%.

**Figure 1 – figure supplement 2.**
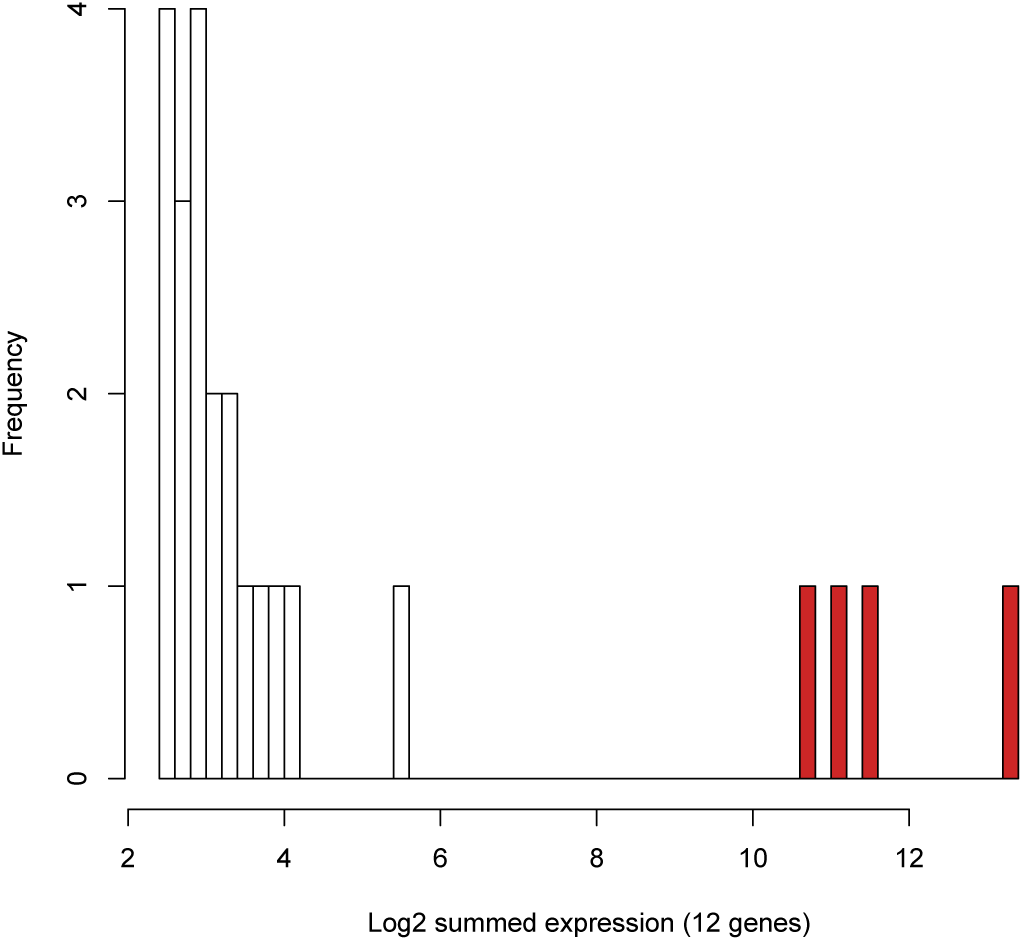
Identification of libraries with female contamination. We summed the expression of the nine chorion genes (CP15 to 19, CP36, CP38) and three yolk proteins (YP1 to 3). We excluded four outlier libraries (in red) with > 256 counts per million bp for the 12 indicator genes.

**Figure 2 – figure supplement 1.**
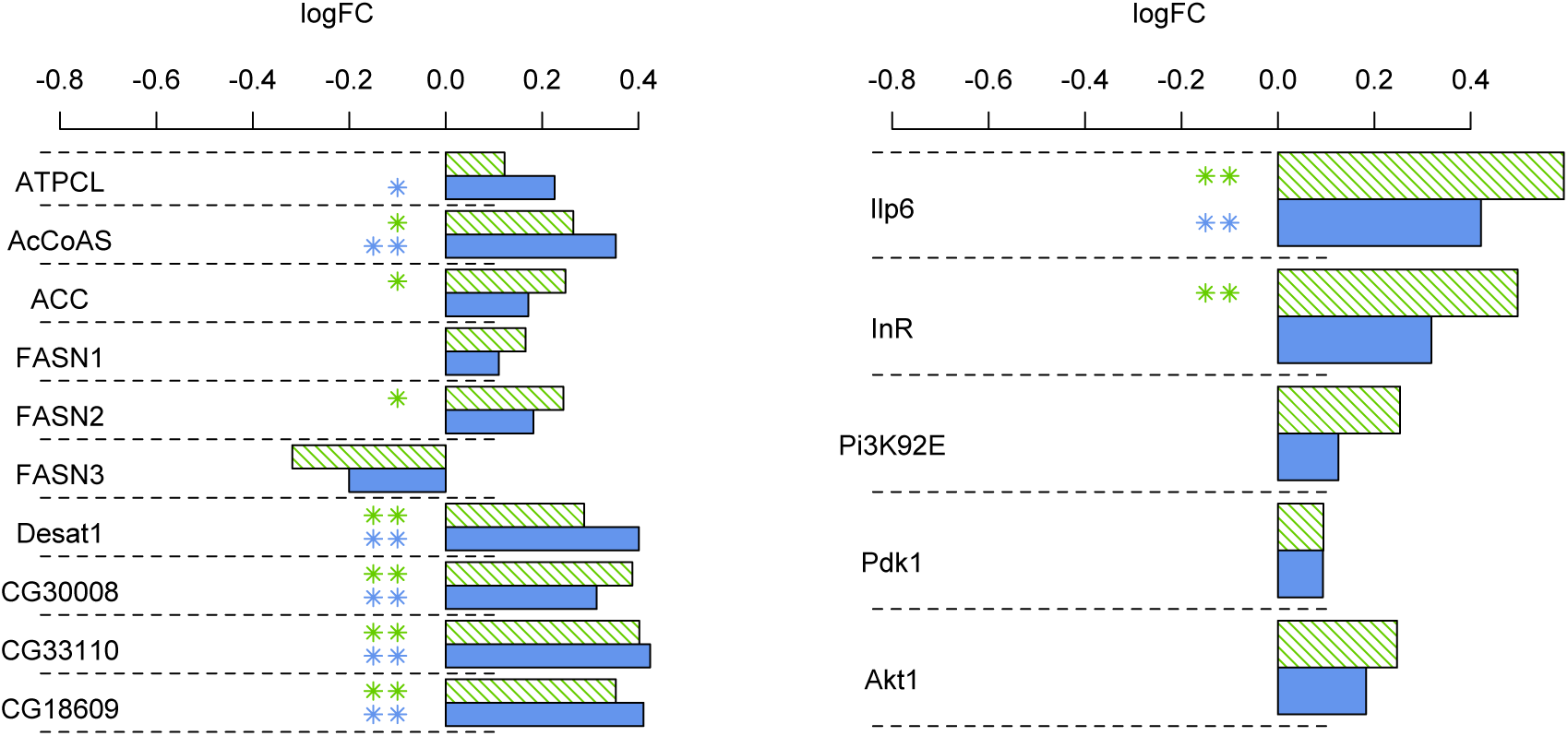
Barplots showing log2 fold change of expression between the hot evolved populations relative to the ancestral (green) or cold evolved (blue) populations. (** FDR<0.05, * FDR<0.1). Left panel: Fat metabolism genes. Right panel: Insulin signaling pathway genes.

**Figure 2 – figure supplement 2.**
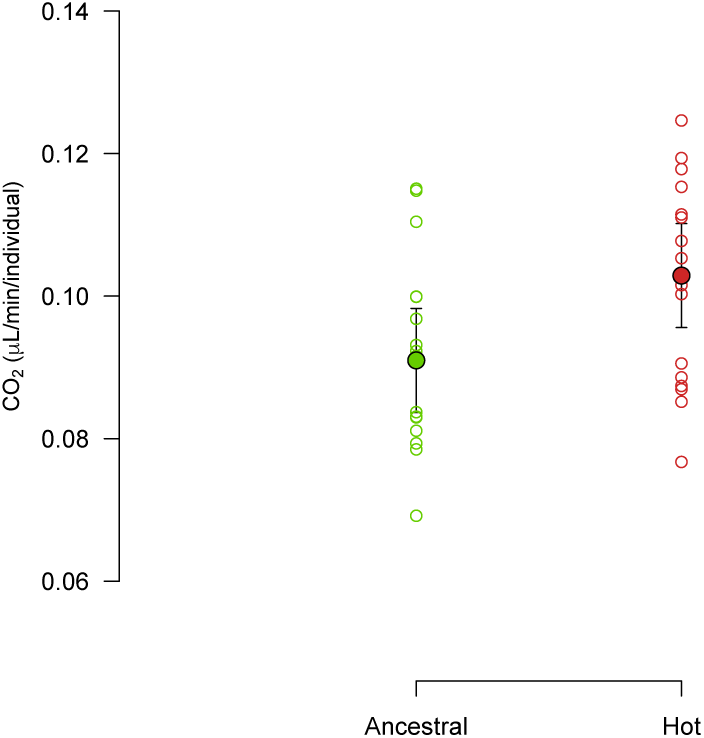
CO_2_ emission is measured over two subsequent generations at 23°C. Although variable, the two mean emission is statistically different between the two populations when the variation in body weight between samples is accounted for. Closed symbols: mean with 95% confidence interval; open symbols: single measurements (n= 2*16), green: reconstituted base, red: population evolved in the hot environment for 133 generations.

**Figure 3 – figure supplement 1.**
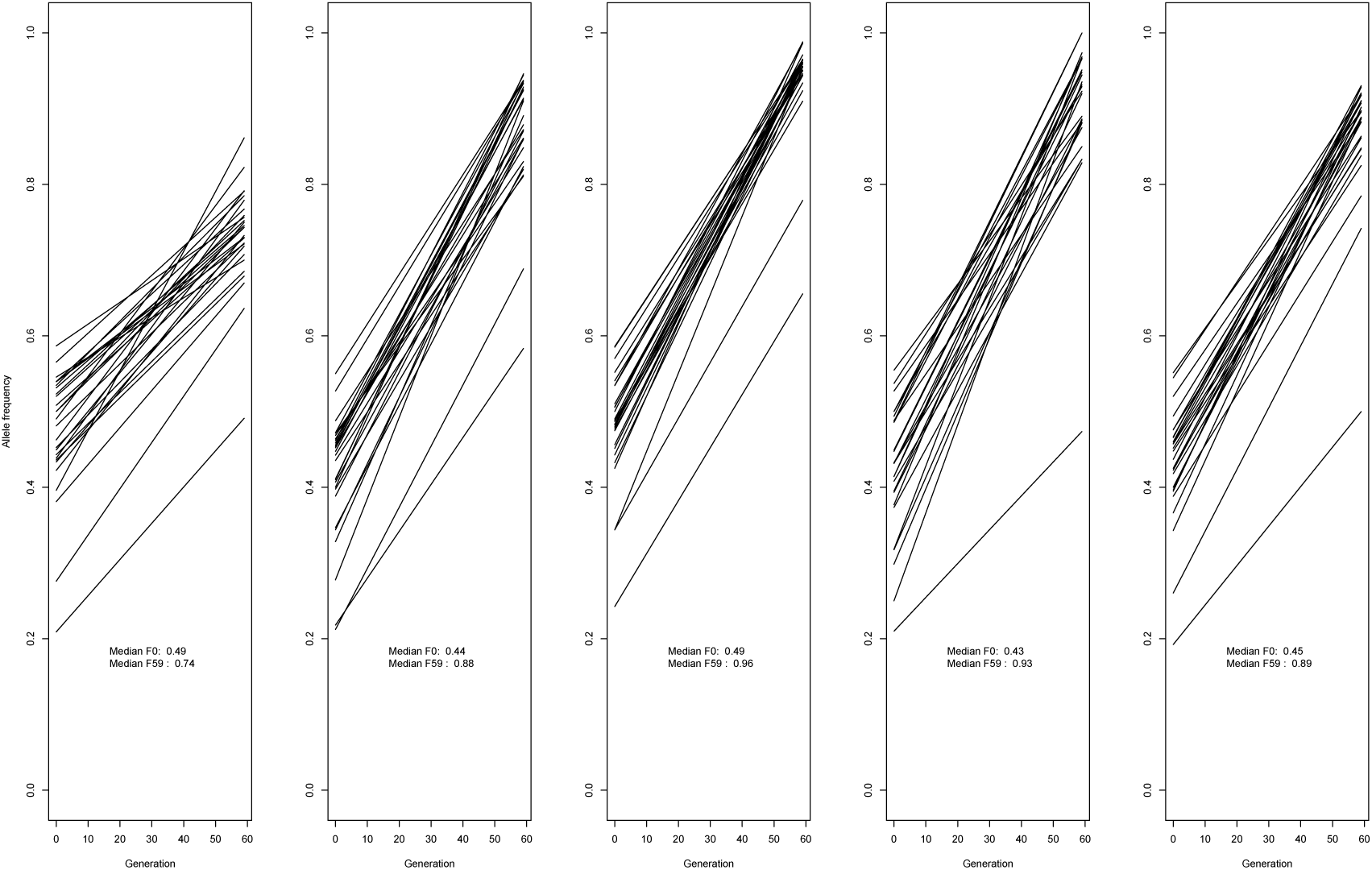
Allele frequency changes during 59 generations of experimental evolution for the 27 SNPs showing a correlated response across 5 hot evolved populations in the *SNF4Aγ* region.

**Figure 3 – figure supplement 2.**
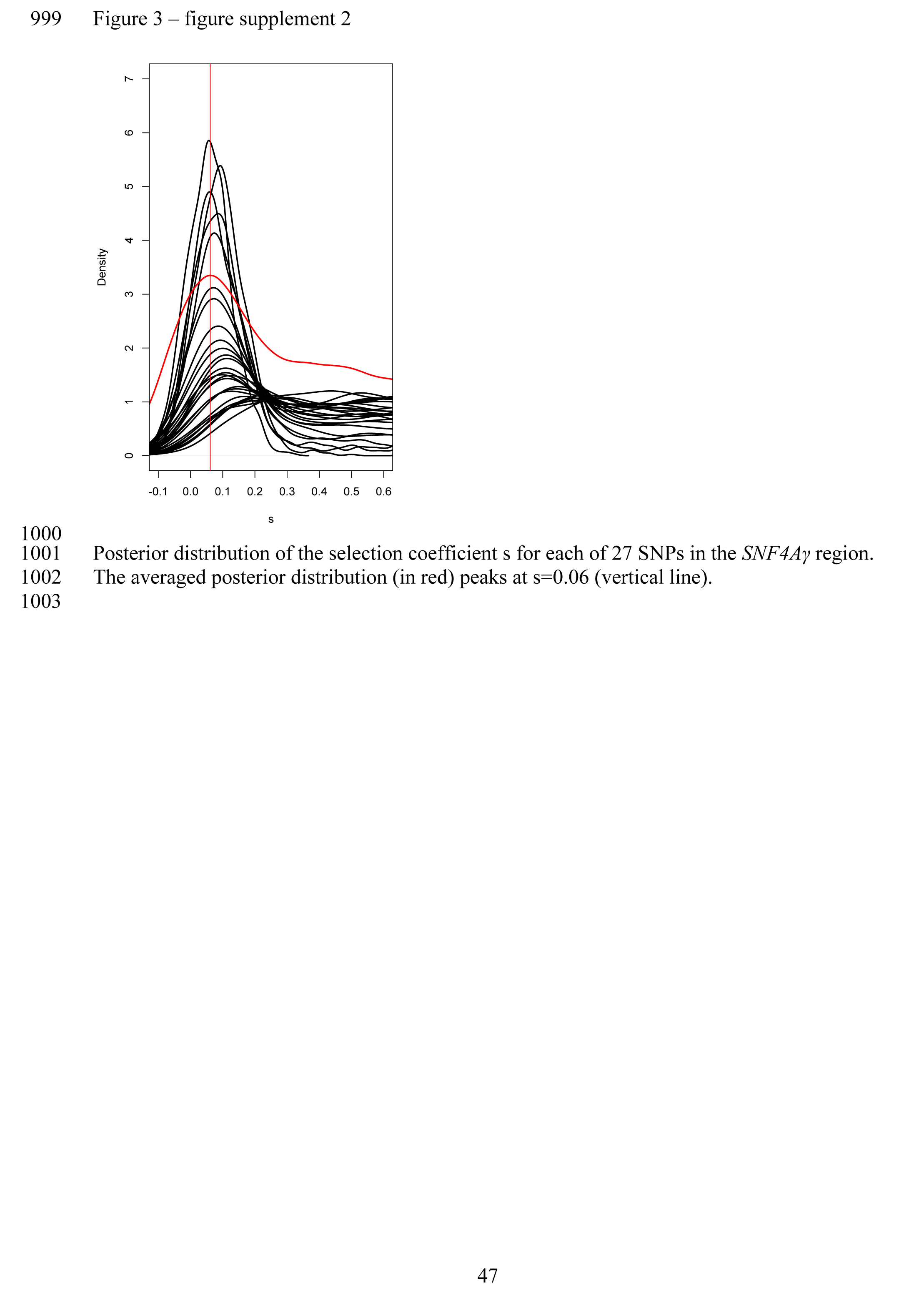
Posterior distribution of the selection coefficient s for each of 27 SNPs in the *SNF4Aγ* region. The averaged posterior distribution (in red) peaks at s=0.06 (vertical line).

**Figure 3 – figure supplement 3.**
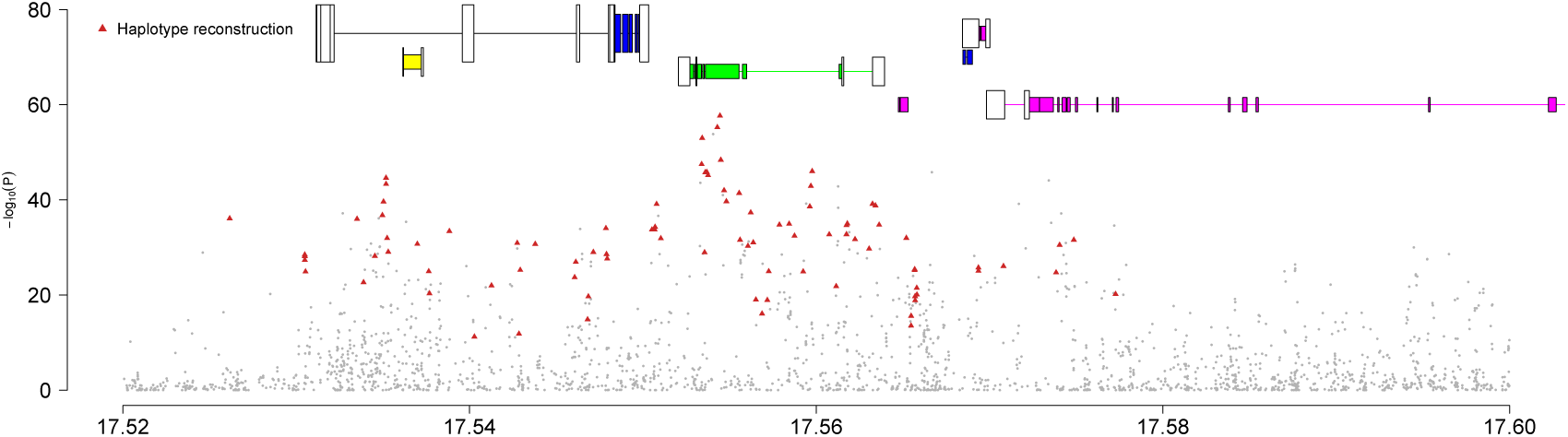
A close up of Manhattan plot around the *Sestrin* region. On top of the Manhattan plot the gene structure of *Sestrin* (leftmost blue gene) is shown. Exons are indicated by colored boxes and introns by thin lines. White boxes indicate UTRs. From left to right the genes are: *Sestrin* (blue), CG18128 (yellow), egl (green), CG13560 (purple), CG11300 (blue), CG5532 (small purple), CG9850 (long purple). Red triangles indicate 96 correlated SNPs, which characterize the selected haplotype(s).

**Figure 3 – figure supplement 4.**
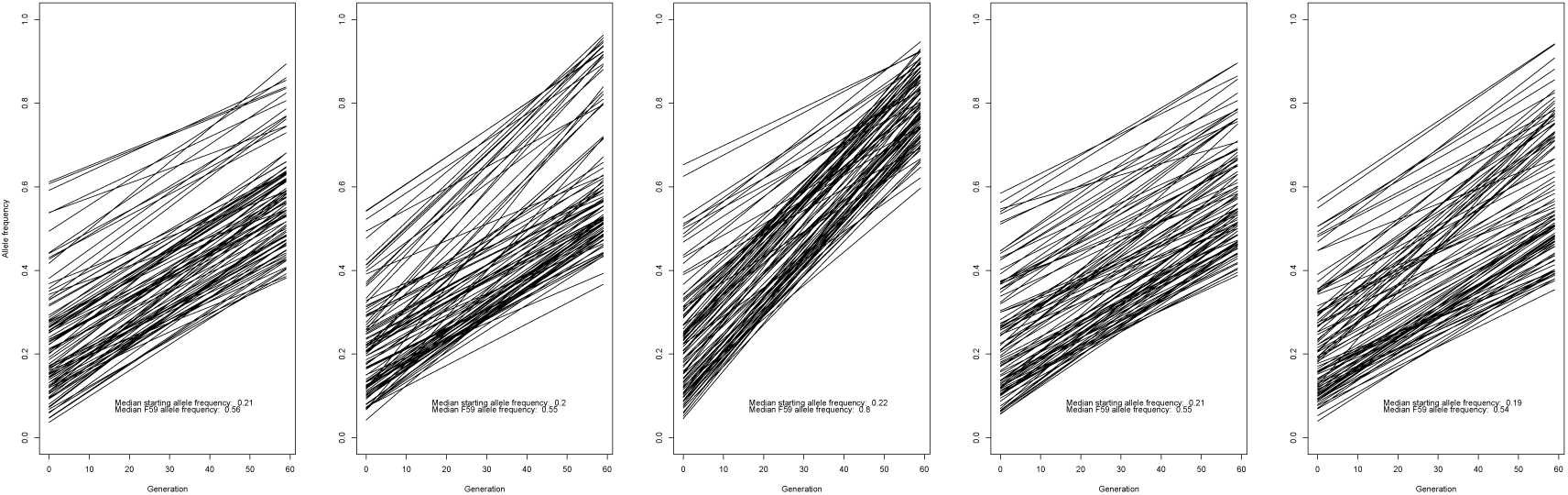
Allele frequency changes during 59 generations of experimental evolution for the 96 SNPs showing a correlated response across 5 hot evolved populations in the *Sestrin* region (2R: 17520000-17600000).

**Figure 3 – figure supplement 5.**
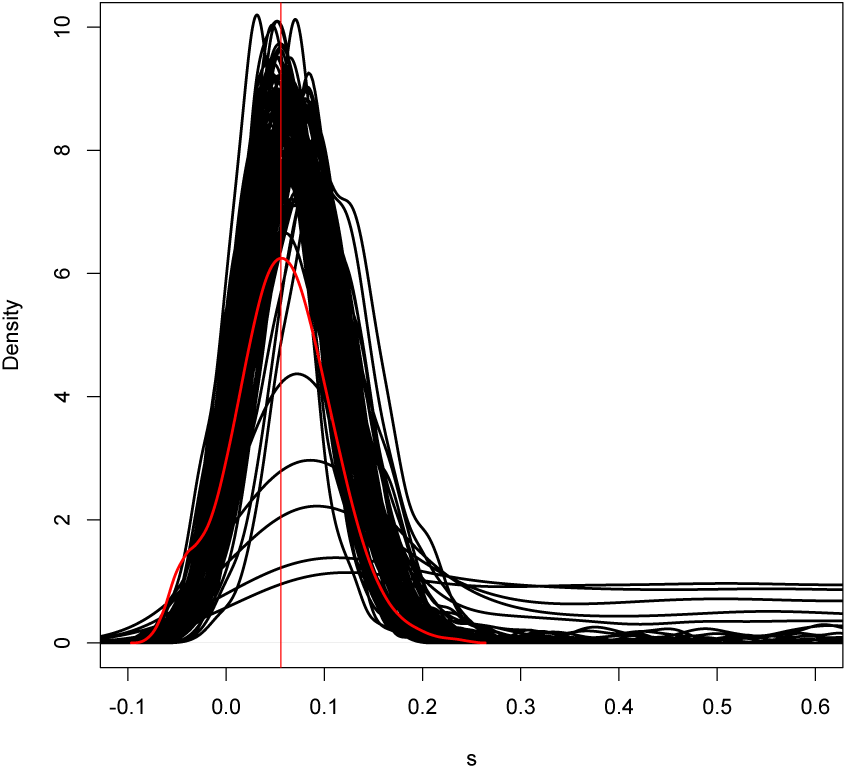
Posterior distribution of the selection coefficient s for each of our 96 SNPs in the *Sestrin* region. The summed posterior distribution (in red) peaks at s=0.055 (vertical line).

**Figure 4 – figure supplement 1.**
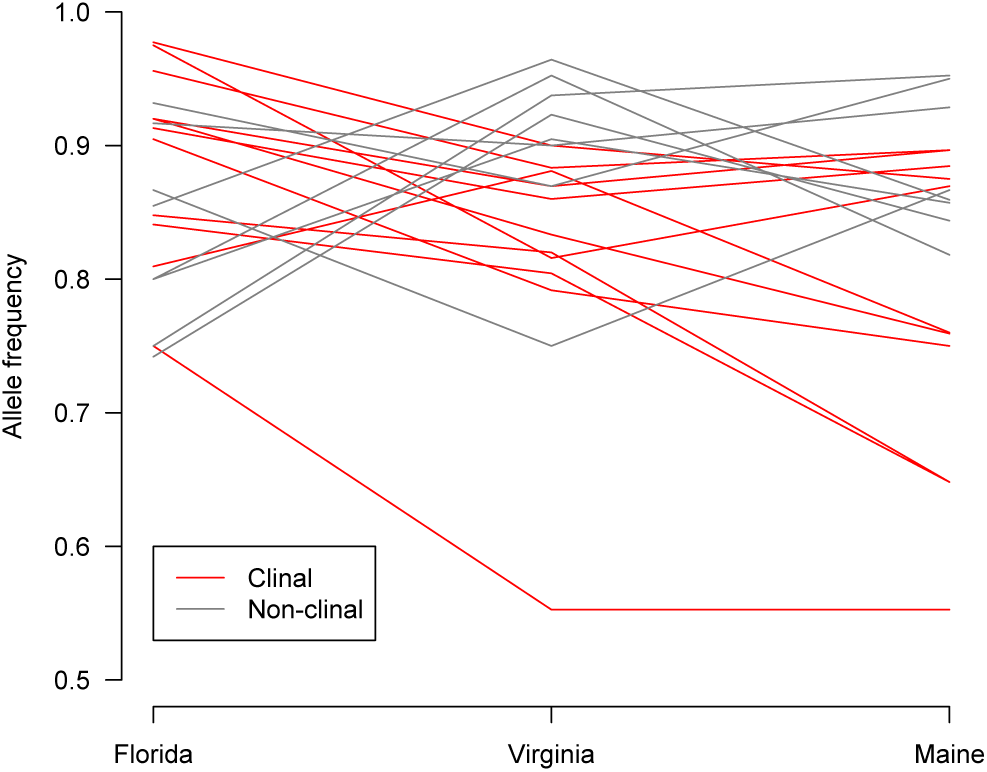
SNPs characterizing the selected haplotype at the *SNF4Aγ* locus in the experimental evolution study show clinal variation in natural populations (Machado et al.). 20 out of 27 SNPs had sufficient coverage in the Machado et al. data set. 11 SNPs displayed clinal variation. The decrease in frequency from Florida to Maine, is consistent with the observation that these alleles are favored in the hot experimental evolution cage.

**Figure 4 – figure supplement 2.**
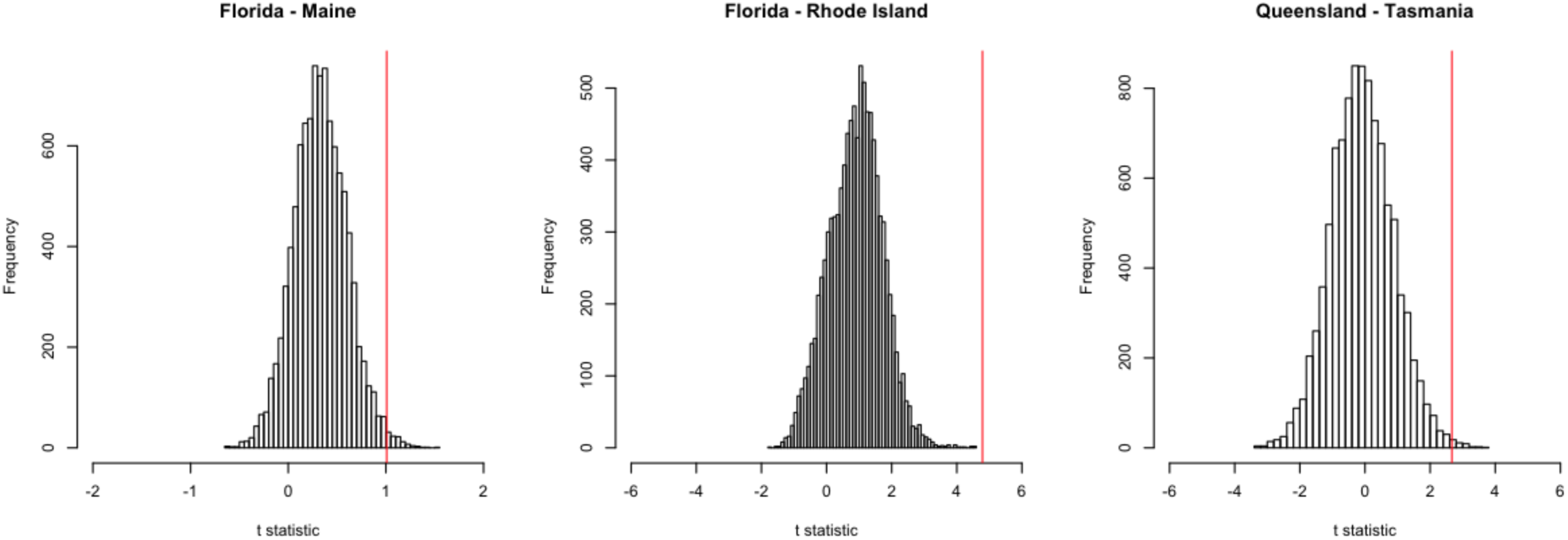
Distribution of t.statistics obtained from random sampling of SNPs in the region of interest for *SNF4Aγ*. In more than 95% of the time the true set of SNPs (red vertical line) are more clinal than randomly sampled SNPs. A. Machado et al. data set (20 SNPs). B & C. Sedghifar et al. data set for B. North American cline (16 SNPs) and C. Australian cline (20 SNPs)

